# Synergy between Membrane Topography and Domains to Control Signaling Protein Localization in Mast Cells Facilitates their Activation

**DOI:** 10.1101/2024.11.22.624791

**Authors:** Shirsendu Ghosh, Alice Wagenknecht-Wiesner, Shriya Desai, Jada Vyphuis, Mariena Silvestry Ramos, John L. Grazul, Barbara A. Baird

## Abstract

Similar to T cells and B cells, mast cell surfaces are dominated by microvilli, and like these other immune cells we showed with microvillar cartography (MVC) that key signaling proteins for RBL mast cells localize to these topographical features. Although stabilization of ordered lipid nanodomains around antigen-crosslinked IgE-FcεRI is known to facilitate necessary coupling with Lyn tyrosine kinase to initiate transmembrane signaling in these mast cells, the relationship of ordered-lipid nanodomains to membrane topography had not been determined. With nanoscale resolution provided by MVC, SEM and co-localization probability (CP) analysis, we found that FcεRI and Lyn kinase are positioned primarily on the microvilli of resting mast cells in separate nano-assemblies. Upon antigen-activation, FcεRI and Lyn merge into overlapping populations together with the LAT scaffold protein, accompanied by merger of microvilli into ridge-like ruffles. With selective lipid probes, we further found that ordered-lipid nanodomains preferentially occupy microvillar membranes, contrasting with localization of disordered lipids to flatter regions. With this proximity of signaling proteins and ordered lipid nanodomains in microvilli, the mast cells are poised to respond sensitively and efficiently to antigen but only in the presence of this stimulus. Use of a short chain ceramide to disrupt ordered-lipid regions of the plasma membrane and evaluation with MVC, CP, and flow cytometry provided strong evidence that the microvillar selective localization of signaling proteins and membrane environments is facilitated by the interplay between ordered-lipid nanodomains and actin attachment proteins, ERM (ezrin, radixin, moesin) and cofilin.

**Significance Statement:** Participation of ordered-lipid nanodomains (aka "rafts") to target immune signaling in the plasma membrane has been established. Separately, membrane topography, specifically microvilli, has also emerged as a participant. Here, we show how these features are coordinated in mast cells that serve as gatekeepers for antigen-triggered, receptor-mediated immune responses, including allergies and inflammation. We found that these specific antigen receptors and a key kinase, together with ordered-lipid nanodomains, localize to microvilli in resting cells, forming separated nano-assemblies. Antigen causes the merger of microvilli into ruffles where receptors and kinases couple to initiate transmembrane signaling. Selective pre-organization of signaling proteins and targeting lipid domains in microvilli and their coordinated redistribution upon antigen stimulation facilitates sensitive and efficient immune responses.

## Introduction

Although cells are often depicted as smooth spheres, microscopic images reveal that the membrane surface of live mammalian cells is irregular, including different types of protrusions. The mechanistic role of this dynamic topography of plasma membranes remains a subject of investigation, with recent studies demonstrating its importance in various fields of cellular physiology (1), including immunology (2–8). Finger-like, actin-dependent microvilli have been found to be the predominant protruding structures on T cells and B cells, which are lymphocytes involved in adaptive immune responses (2–6, 8–10). To elucidate participation of microvilli in immune cell function the topographical distribution of proteins and lipids must be quantitatively assessed. This has been technically challenging because the width of microvilli lies within ∼100 nm, below the diffraction limit of the optical microscope.

Recently, Haran and co-workers designed a super-resolution microscopy-based methodology called Microvillar Cartography (MVC) to examine the localization of membrane proteins with respect to the plasma membrane topography in fixed samples (2, 4, 5, 11). They found strong evidence that the microvilli of Jurkat and human effector T cells serve as signaling hubs for T cell receptor (TCR)-mediated responses to antigen in cellular immune responses. They observed that TCR, co-receptor CD4, tyrosine kinase Lck, and adaptor LAT are substantially enriched in the microvilli of resting cells. This pre-assembly can thereby facilitate initial antigen sensing such that TCR couples effectively with its proximal signaling proteins. Consequent cell activation then modulates membrane linkages with filamentous actin mediated by ezrin-radixin-moesin (ERM) proteins (5). Consistent with this view, Jun and co-workers showed that TCRs are on the tips of microvilli (12) and that microvilli-derived particles can form “immunological synaptosomes” with antigen-presenting cells (APCs) (6). Krummel and coworkers determined that microvilli serve as sensors for T-cells for finding specific APCs (3). In other studies, Reth and coworkers demonstrated that localization of antigen-sensing B cell receptors (BCRs) is similarly dictated by B cell microvilli (8).

All these studies point strongly to microvillar-mediated signal initiation as a key mechanism for immune cells, as represented by T- and B-lymphocytes. Mast cells play different roles in innate and adaptive immunity, notably in allergic and inflammatory responses (13). For mast cells, the high-affinity receptor (FcεRI) for immunoglobulin E (IgE) is the dominant antigen-sensing receptor that is mediated by tightly bound, antigen-specific IgE (14). A microvillar topography like T cells and B cells has been characterized for primary mast cells (15–18) and for Rat Basophilic Leukemia (RBL) mast cells (19). However, the relationship between FcεRI, signaling partners, and the microvilli has not been defined.

Extensive investigations of the spatial relationship of antigen receptors and their signaling partners in the flattened ventral membrane of attached immune cells has been carried out using TIRF microscopy and quantification of super resolution images. For T cells (20), B cells (21), and mast cells (22), antigen engagement of their respective receptors causes physical and functional coupling with key tyrosine kinases, resulting in phosphorylation of the immune receptors, assembly of additional proteins and consequent downstream signaling. The targeting role played by membrane lipid heterogeneity has been a subject of considerable interest (23). With lipid compositions of model membranes representing those in the plasma membrane, nanoscopic regions of ordered lipids (characterized by saturated acyl chains and cholesterol) segregate from regions of disordered lipids, described as Lo-like and Ld-like, respectively. Although ordered-lipid nanodomains in cellular plasma membranes are diverse in lipid and protein composition, these are often lumped loosely together and colloquially dubbed “rafts.” Many different studies have examined participation of “rafts” or Lo-like ordered-lipid nanodomains to initiate signaling in mast cells, T cells and B cells (21, 24–27). Compelling evidence supports the view that this phase-like behavior of lipids is important for regulating protein interactions in the membrane, and these may also couple to phase-based protein condensates in the cytoplasm (28–30) .

Participation of ordered-lipid nanodomains to regulate protein distributions during antigen-activation, in the context of plasma membrane topography, has been largely unexamined in any immune cell type. For the present study, we adopted the MVC methodology and utilized RBL mast cells as a model system (31) to spatially relate membrane nanodomains to microvilli and evaluate their coordination in targeting antigen-stimulated mast cell signaling. We quantified the distribution of FcεRI and key signaling proteins, as well as order- and disorder-preferring lipid probes with respect to the plasma membrane topography of resting suspended cells. We observed how these protein and lipid distributions change after antigen is introduced to crosslink FcεRI, causing cell activation. Like T cells examined in previous studies (2, 4, 5), we found that antigen receptor and key signaling kinase localize to microvillar tips in RBL cells, but that the stimulated morphological changes differ markedly from those of T cells. We also investigated the participation of cytoskeleton attachment proteins, ERM and cofilin, as have been reported to regulate localization of signaling proteins in immune cells (5, 32, 33). Our study uncovers the complex interplay between membrane topography, ordered-lipid nanodomains, and cytoskeletal attachment proteins in controlling the distribution of signaling proteins, which together sense the presence of antigen and initiates cell activation such as occurs during immune responses.

## Results

### Activation of mast cells is accompanied by changes in topographical features

The topography of RBL mast cells (19) resembles primary mast cells (15–17) as well as T- and B-lymphocytes (2, 4, 5, 9, 34, 35), which play complementary roles in immune responses. We employed RBL cells (RBL-2H3 cell line) in the present study as a robust representative for mast cells more generally (31, 36). The surfaces of resting RBL cells are dominated by microvilli (Fig. 1A). However, unlike activation and synapse formation of T-cells (37), SEM shows that activation of mast cells by soluble antigen does not cause immediate microvillar collapse (18, 19). RBL cells were sensitized to antigen (DNP-BSA) by saturating FcεRI with anti-DNP IgE. We found that short activation (1 min) of suspended RBL cells with soluble antigen causes the microvilli to merge (Fig. 1B). Prolonged activation (15 min) results in the formation of long and curvy ridges, commonly called “ruffles” (Fig. 1C). Our observations with SEM are like those made previously with adherent RBL mast cells (19), and confocal microscopy of live RBL cells shows the ruffles to be dynamic (38, 39).

**Figure 1.**
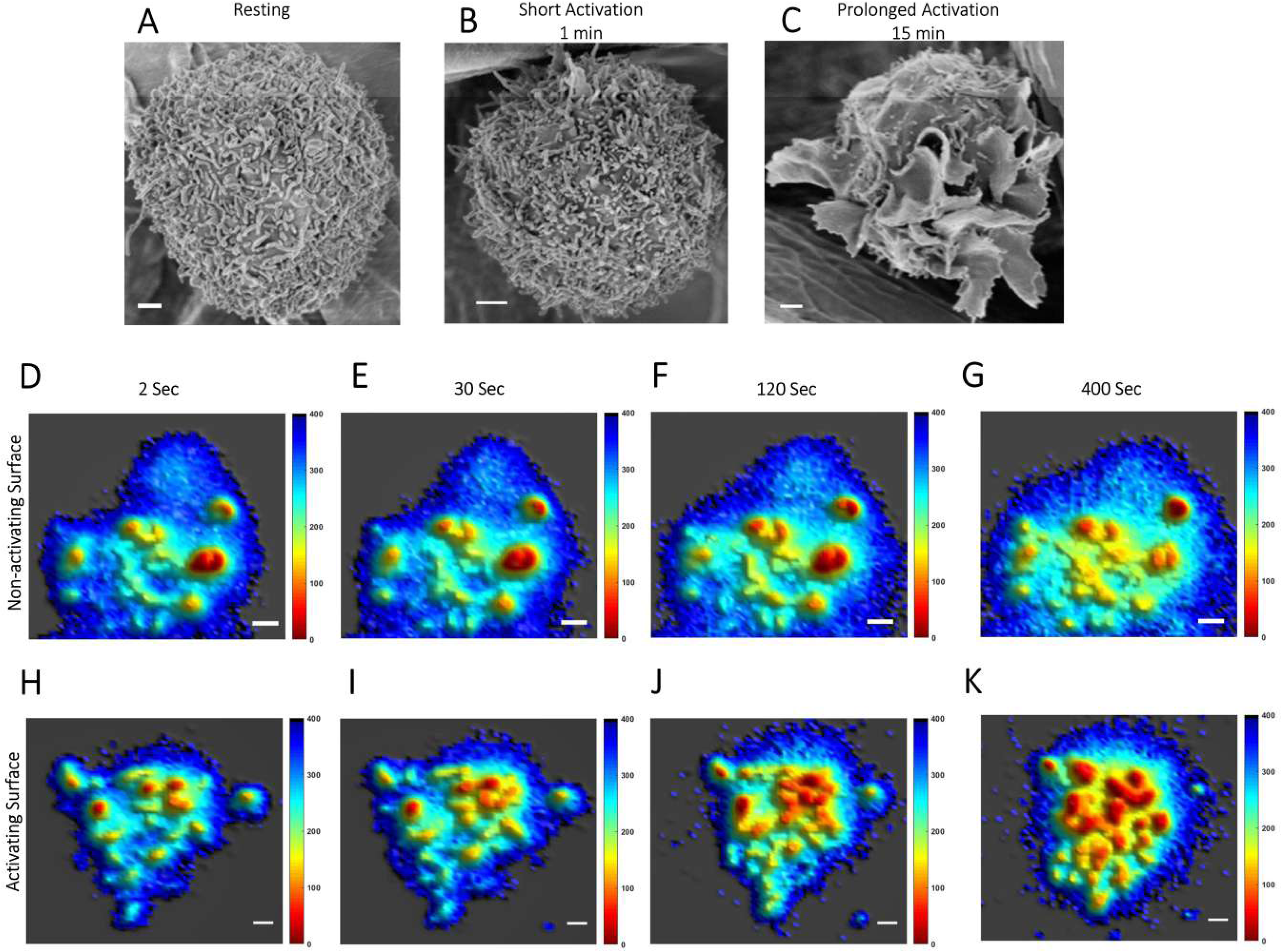
Membrane Topography of RBL Mast Cells is Dominated by Microvilli which form Membrane Ruffles upon Activation: A-C. SEM images of RBL cells. A. Non-sensitized, suspended resting cells. B-C. After 1 minute (B) or 15 minutes (C) activation of suspended anti-DNP IgE-sensitized RBL cells using soluble antigen (DNP-BSA). Scale bar, 1μm. D-K. Snapshots from movie of membrane topographical change of IgE-sensitized RBL cells while interacting with BSA coated non-activating glass surface (D-G) and 20 mol% DNP-BSA/BSA coated activating glass surface (H-K). Complete movies are provided in SI Appendix Movies S1 and S2. Scale bar, 1μm.

We adopted MVC methodology (11) (SI Appendix Fig. S1) to further delineate the changes in membrane topography and the distributions of FcεRI and key signaling proteins before and after activation by antigen. MVC evaluates the plasma membranes of cells uniformly labeled fluorescently with dyes FM143 (live cells) or FM143fx (fixed cells) (5, 40, 41), and utilizes variable-angle total internal reflection fluorescence microscopy (VA-TIRFM) to quantify membrane topography with 20 nm resolution in the z direction (42). The relative distance of each point on a TIRFM image from the glass surface (δz) is determined from the measured fluorescence intensity at that point for each of several selected angles of incidence of the excitation beam (SI Fig. S1D-F) (5, 11, 43). The 3D-membrane topography map is then created from the set of δz values obtained by combining all the VA-TIRFM images (SI Fig. S1G,H), which increases the statistical precision. Because the typical width of a microvillus (∼100 nm) is below the optical diffraction limit, the microvillar structure appears thicker (∼ 250 nm diameter) in X-Y direction of VA-TIRFM images, which can achieve sub-diffraction resolution only in the Z-direction.

In one set of experiments, suspended live RBL cells were sensitized with anti-DNP IgE and stained with FM143 (5). Then the cells were dropped on glass surfaces coated with either BSA (non-activating) or BSA containing 20 mol% DNP-BSA (antigen-activating). We recorded a time series of VA-TIRFM images of cells interacting with these two different surfaces using MVC methodology (10, 11). Five independent movies of cells under each of these two conditions were recorded, with each cell imaged from an independent experiment. Post-processing of these live cell MVC image-time series allows the dynamics of membrane topography during the interactions to be observed, and representatives are shown in SI Appendix Movies S1 and S2. We found that for cells interacting with the non-activating surface the membrane topography of mast cells remains almost unaltered for up to ∼400 seconds (Fig. 1D-G; SI Movies S1A and B). Within 30 min the membrane begins to flatten and adhere to the glass surface as expected from previous studies showing integrin and cytoskeleton engagement (39, 44). In contrast, microvilli on cells interacting with the antigen-activating surface start to merge within 30 sec of interactions (Fig. 1H-K; SI Movies S2A and B). Laterally extended structures (as long as 3 μm) are observed after several minutes of activation (Fig. 1J,K). The longest dimension of a microvillus, whether projecting straight up from the cell surface or bent over, typically ranges 0.1 μm - 1 μm, with a median length around 0.3 to 0.4 μm. In contrast, ridge-like ruffles, are typically longer than 1 μm and can extend several microns (35). Accordingly, we classified the long projections that develop during activation as ruffles, consistent with the view provided by SEM (Fig. 1C). As seen dynamically in SI Movie S2A, these ruffles form through the merging of microvilli during prolonged interaction with the activating surface (Fig. 1H-K). At longer times the cells flatten and spread as observed previously in studies with activating surfaces (39, 44).

### Membrane topography and localization of signaling molecules are correlated

We used MVC on fixed RBL mast cells to examine the spatial relationship between membrane topography and membrane proteins that are involved in transmembrane signaling after immune stimulation. Crosslinking of IgE-FcεRI complexes by antigen causes coupling with Lyn tyrosine kinase, which is anchored to the inner leaflet of the plasma membrane and phosphorylates cytoplasmic segments of FcεRI. Consequent recruitment of cytoplasmic Syk tyrosine kinase leads to phosphorylation of transmembrane LAT, which serves as a scaffold for downstream signaling proteins. To investigate distributions of FcεRI, Lyn, and LAT we fixed suspended cells, either non-sensitized (resting) or IgE(anti-DNP)-sensitized in a resting state or after activation for 1 min or 15 min with soluble antigen (DNP-BSA) at 37°C. For Lyn or LAT, cell membranes were homogeneously stained with FM143fx, followed by fixing and permeabilizing these cells, and labeling with the respective Alexa- 647 tagged antibody. FcεRI was labeled by Alexa-647 tagged IgE during overnight cell culture, and this was followed by fixation and membrane staining with FM143fx. As we observe routinely, overnight sensitization of RBL cells with monomeric IgE does not cause cell activation (45). The antibody-labeled, membrane-stained, fixed suspended cells were attached to a poly-D-lysine (PDL) coated glass surface and imaged with TIRF microscopy. Because the cells are pre-fixed, this surface interaction causes no perturbation of the topography or activation of the cells (5). We observed similar cell topography in our MVC images (microvillar density = 3.8±0.5/μm^2^, calculated from 10 MVC images of resting cells) as in our SEM images (microvillar density = 4.4±0.5/μm^2^, calculated from 5 SEM images of resting cells), confirming the robustness of the MVC methodology. Similar microvillar density (2-4/μm^2^) was reported for T-cells (35).

In the next step, we mapped the locations of the fluorescent antibody-labeled membrane proteins in the X-Y direction with accuracy below the diffraction limit using stochastic localization nanoscopy (SLN) (SI Fig. S1B). This provides the nanoscale distribution pattern (in X-Y direction) of the specifically labeled membrane protein in the context of nanoscale membrane topography (in Z direction; SI Fig. S1A) by superimposing the SLN map of membrane proteins on the VA-TIRFM membrane topography map (SI Fig. S1C). Because we are concerned only about the distribution of the membrane proteins (and not their absolute number), multiple fluorophores per labeling antibody do not lead to an artifactual conclusion. We recorded the SLN-based protein distribution map in two separate slices for each cell (SI Fig. S1K). One slice is at the glass surface (0 nm), typically visualizing mainly the microvillar region and a small fraction of the cell body region (for shorter or bent-over microvilli). The other map is at a slice 400 nm away from the glass surface where both microvilli and cell body regions are included for most microvilli. We demonstrated previously that the TIRF evanescent field intensity, although decreasing with distance from the glass surface, is still high enough to detect fluorophores present in 0 nm and 400 nm slices with equal probability (5).

The membrane topography map of each cell was segmented into either microvillar region or cell-body region following the established MVC methodology (11). From our combined measurements we quantified four features in each cell: 1) the percentage of the total microvillar area relative to the total cell area, as projected in 0 nm and 400 nm imaging slices (SI Fig. S1I,J); 2) the fraction of labeled proteins of each type residing in the microvilli in 0 nm and 400 nm imaging slices (SI Fig. S1I,J); 3) the fraction of total microvilli occupied by each labeled protein; 4) the fluorophore density (δCount/δArea) as a function of radial distance (R) from the tips of microvillar structures in the 0 nm imaging slice (SI Fig. S1L-N). The central microvillar region is defined as the region within 20 nm in the Z-direction from the pixel with the minimum δz value (microvillar tip).

Importantly, the topographical analysis relies only on the sub-diffraction resolution in the Z-direction, as determined by VA-TIRFM. MVC-based superposition of topography and SLN allowed us to determine the distribution of endogenous membrane proteins involved in FcεRI-mediated signaling in RBL mast cells: FcεRI, Lyn kinase, and LAT scaffold. We also investigated ectopically expressed, transmembrane tyrosine phosphatase, PTPα, which counteracts Lyn kinase activity (46). We made these measurements for suspended cells in the resting state and after 1 min and 15 min activation with soluble antigen. A summary of our findings is described below (Fig. 2 and SI Fig. S2).

**Figure 2.**
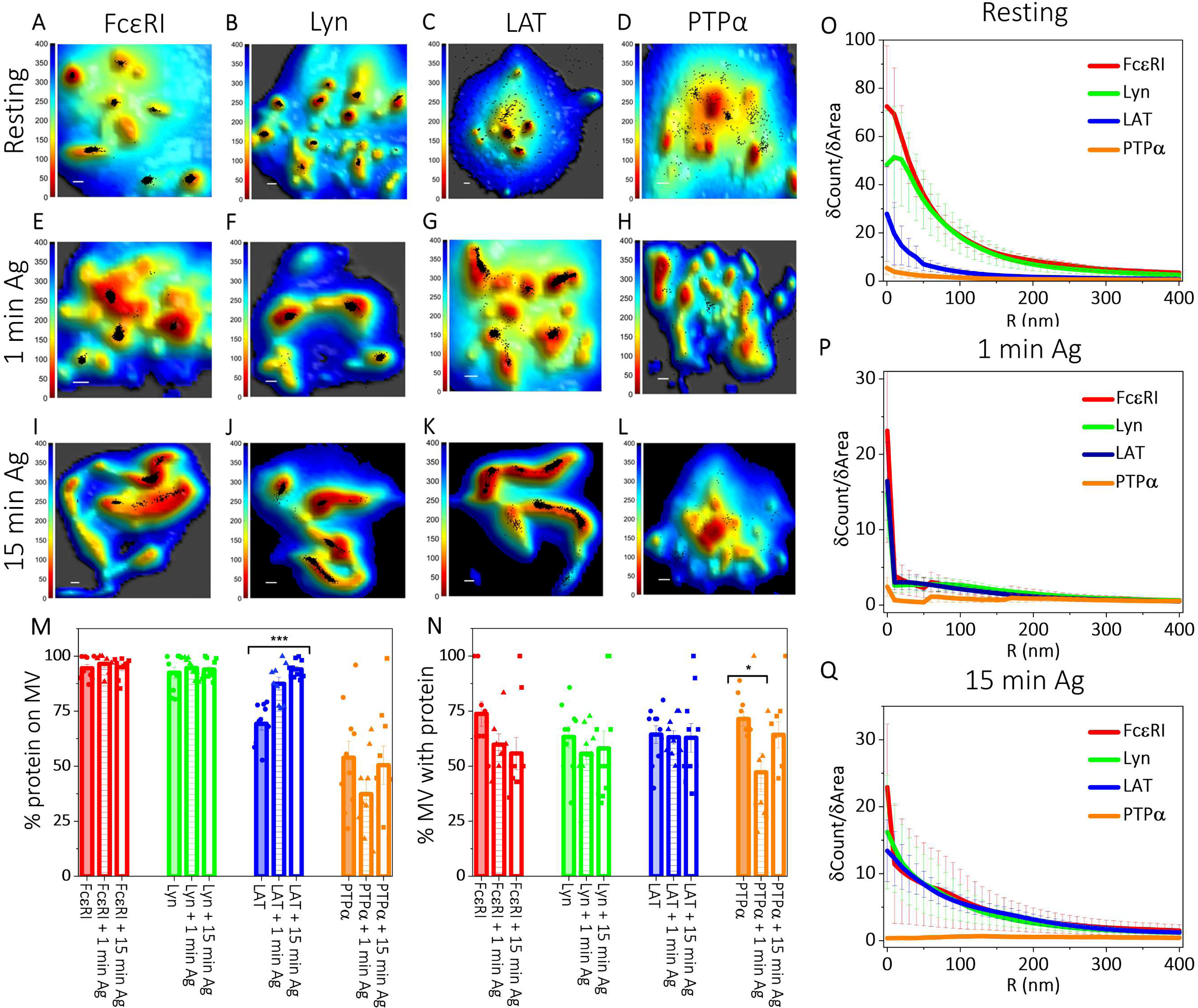
Membrane Topography and Localization of Signaling Proteins are Correlated in Resting and Antigen-Activated States of RBL Mast Cells: A-L. Localization maps of RBL cell membrane signaling proteins with respect to the 3D membrane topography for non-sensitized suspended resting cells (A-D), and for sensitized cells after 1 min (E-H) or 15 min (I-L) activation with soluble antigen. Positions of proteins obtained from SLN (black dots) in the 0 nm imaging slice are superimposed on membrane topography maps obtained from VA-TIRFM. The color bars represent the z distance from the glass in nanometers (nm). Scale bars (x,y), 0.5 μm. M-Q. Quantified distributions of proteins on the RBL cell surface in resting, 1 min and 15 min antigen-activated states. The error bar represents the standard error of the mean. M. Percentage of specified proteins on microvillar (MV) regions of the membrane in resting, 1 min, and 15 min activated states in the 0 nm slice. The values for individual cells are shown as dots in the plot. N. Fraction of microvilli (MV) occupied by the specified proteins in resting, 1 min and 15 min activated states. The values for individual cells are shown as dots in the plot. O-Q. Protein fluorophore density (δCount/δArea) as a function of radial distance (R) from the central microvilli region in the 0 nm imaging slice in resting (O), 1 min (P), and15 min (Q) activated states. P-values for M and N are given in SI Appendix Table S1

#### FcεRI

FcεRI is specifically labeled with monomeric IgE. Crosslinking of FcεRI complexes by soluble antigen causes the formation of nanoclusters to activate the cell (26, 27, 30).

##### Resting state

We observed that 94±2% of FcεRI localize to microvilli in the 0 nm slice (Fig. 2A,M). The pattern is very similar in the 400 nm slice, 92±3% FcεRI localize to microvilli (SI Fig. S2E,Q). This evaluation by MVC is consistent with TIRF images of these cells labelled with Alexa-647 tagged IgE, which show fluorescent puncta as expected for IgE-FcεRI localized to microvilli (SI Fig. S2A). 74±6% of microvilli are occupied by FcεRI (Fig. 2N). The density plot for FcεRI (δCount/δArea *vs* R) decays sharply in the 0 nm SLN slice as the radial distance (R) from the central microvillar region increases, further indicating predominant localization nearer the microvillar tips (Fig. 2O).

##### Changes with activation

Incubation with soluble antigen for 1 min or 15 min causes no significant changes in localization of FcεRI with respect to the microvilli although these structures change from separate microvilli to merged microvilli and ruffles during this time period (0 nm slice: Fig. 2A,E,I,M; 400 nm slice: SI Fig. S2E,I,M,Q). Quantitatively, the percentage of microvillar area relative to total cell area increases significantly from 15±2% in resting cells to 23±2% after short (1 min) antigen-activation of IgE-sensitized cells and further rises to 37±2% after prolonged activation (15 min) of these cells (SI Fig. S2R). The fraction of microvilli occupied with FcεRI appears to decrease slightly upon activation (Fig. 2N), although the decrease is not statistically significant (P>0.05, SI Table S1). The δCount/δArea *vs* R plot for FcεRI becomes significantly sharper upon activation for 1 min, consistent with the view that more FcεRI are accumulating towards the tips of the microvilli (Fig. 2P). By 15 min activation the δCount/δArea plot becomes more gradual, probably reflecting the formation of ruffles rather than sharp microvilli (Fig. 2Q).

#### Lyn

Cytoplasmic, membrane-anchored Lyn couples with and phosphorylates antigen-clustered FcεRI as the first step of transmembrane signaling (22, 27). This coupling involves partitioning of Lyn, via saturated palmitate and myristoylate chains, into ordered-lipid nanodomains that coalesce around the clustered FcεRI (27, 47, 48) .

##### Resting state

Like FcεRI, most Lyn localizes in the microvilli of RBL cells in imaging slices at 0 nm (92±3 %; Fig. 2B,M) and 400 nm (91±2 %; SI Fig. S2F,Q). 63±5 % of microvilli are occupied by Lyn (Fig. 2N). Microvillar localization was also observed in TIRF images of these cells (SI Fig. S2B). Interestingly, the δCount/δArea *vs* R plot for Lyn shows a peak at ∼20 nm away from the central microvillar region then decays sharply (Fig. 2O). This is a distinctive difference from the same plot for FcεRI, which decays sharply directly from the central microvillar region (Fig. 2O), suggesting an inherent separation of FcεRI and Lyn within the same microvillus for resting RBL mast cells. In a subsequent section, we describe co-localization probability (CP) analysis to quantify further the separation of these two proteins within microvilli.

##### Changes with activation

Antigen activation for 1 min or 15 min causes little or no change in the percentage of Lyn on the microvilli (Fig. 2M and SI Fig. S2Q) or the percentage of microvillar structures occupied by Lyn (Fig. 2N). However, antigen activation for 1 min causes the δCount/δArea plot for Lyn kinase to decay steeply from the central microvillar region (Fig. 2P), and this decay appears to become more gradual after 15 min activation (Fig. 2Q). The δCount/δArea plots for Lyn after both 1 min and 15 min activation have nearly the same shape as the corresponding plots for FcεRI, consistent with the view that antigen-clustering of FcεRI causes coupling of these signaling molecules on the microvilli.

#### LAT

LAT is primarily cytoplasmic and anchored to the membrane by a short transmembrane segment; reversible palmitoylation enhances partitioning into ordered-lipid nanodomains (49). After antigen-clustering of FcεRI, palmitoylated LAT serves as a scaffold for the assembly of proteins necessary for downstream signaling (27, 50). LAT plays a similar role in T cells (20), and previous MVC studies showed that ∼70% of LAT molecules reside in the microvilli of resting T cells (5).

##### Resting state

Compared to FcεRI and Lyn, LAT localizes less prominently in the microvilli of RBL cells in imaging slices at 0 nm (69±3%; Fig. 2C,M) and 400 nm (68±3%; SI Fig. S2G,Q). 64±4% of the microvilli of the cells are occupied with LAT molecules (Fig. 2N). TIRF images of these cells also show fewer microvillar puncta and more fluorescence in cell body regions compared to FcεRI and Lyn (SI Fig. S2C). The δCount/δArea *vs* R plot for LAT exhibits a decay shape that is shallower compared to those of FcεRI and Lyn further indicating that although LAT proteins tend to localize to microvilli, they also distribute across the cell body (Fig. 2O).

##### Changes with activation

The percentage of LAT localized in the microvilli in the 0 nm slice increases from 1 min (87±3%) to 15 min (94±1%) after antigen-clustering of FcεRI (Fig. 2G,K,M), and a similar increase is observed in the 400 nm slice (SI Fig. S2K,O,Q). This increase with activation is statistically significant (P << 0.5 SI Table S1). The fraction of microvilli occupied by LAT remains almost unchanged after antigen activation (Fig. 2N). By 1 min the δCount/δArea plot for LAT increases in magnitude at the microvillar central region with a steeper decay and a shape that is very similar to those of FcεRI and Lyn (Fig. 2P). These three decay plots continue to be very similar after 15 min (Fig. 2Q), consistent with the coalescence of these three signaling proteins on microvilli upon antigen activation.

#### PTPα

By counterbalancing tyrosine kinase activities, transmembrane tyrosine phosphatases are known to regulate the activation of immune cells including mast cells (51), such that the spatial relationship of these proteins to microvilli is of considerable interest. CD45 has been established as a negative regulator in T cells, and previous MVC studies showed that this protein distributes uniformly between microvilli and flatter cell regions (5, 52). CD45 appears to have positive and negative regulatory roles in mast cells (53). Moreover, this protein is expressed in low levels in normal mast cells, including RBL cells (46, 54), which precluded MVC-based distribution map of CD45 in these cells with high confidence. Instead, we chose to investigate ectopically expressed PTPα, a transmembrane tyrosine phosphatase that we found previously to negatively regulate Lyn kinase activity in a reconstituted system (46). Our previous studies showed that PTPα localizes preferentially in disordered-lipid regions, away from antigen-clustered FcεRI which stabilizes ordered-lipid nanodomains domains proximally (27).

In MVC studies, we transiently transfected RBL cells with human influenza hemagglutinin (HA)-tagged PTPα. Our transfection efficiency for HA-PTPα was ∼30%, and with Alexa-647 labeled anti-HA antibodies we could identify the transfected population. Transfected cells were fixed, stained with FM143fx, and labeled with fluorescent antibodies after permeabilization. Interestingly, we found that microvilli in these samples depended on the level of transfection as visualized with fluorescence. Although cells with lower levels of PTPα expression looked like the samples stained for FcεRI, Lyn, or LAT, cells with high expression levels of PTPα exhibited much fewer microvilli.

For MVC analysis we selected transfected cells that had moderate levels of PTPα expression. For those cells we found 54±7% and 58±6% of PTPα on the microvilli in 0 nm and 400 nm slices, respectively (Fig. 2D,M; SI Fig. S2H,Q). We found that 72±3% of microvilli of those transfected cells have some PTPα (Fig. 2N), but TIRF images show the fluorescent label spread across the cell body, markedly different from respective of FcεRI and Lyn and LAT (SI Fig. S2D). The slope of δCount/δArea vs R plot for PTPα is very shallow compared to those for FcεRI, Lyn, and LAT, consistent with an almost uniform distribution of PTPα across the membrane microvilli and cell body (Fig. 2O). When these cells transfected with PTPα are incubated with antigen to cluster FcεRI, we found that their membrane topography changes are disrupted compared to those observed for FcεRI, Lyn, and LAT described above: Merger of microvilli and formation of ruffles do not occur in the same manner (Fig. 2D,H,L; SI Fig. S2H,L,P).

To evaluate how HA-PTPα transfection of RBL cells affects their function, we performed an assay of the ultimate cell response to effective stimuli: release of secretory granules (degranulation) (45). We found that the percentage of degranulation stimulated by antigen is decreased by ∼25% for HA-PTPα transfected cell batches compared to cells that had not been transfected (SI Fig. S2S). Comparing this result to a transfection efficiency of ∼30% suggests that our transfection of PTPα substantially inhibits cellular degranulation. Thus, although we imaged the change in the distribution of moderately expressed PTPα before and after antigen-activation of transfected RBL mast cells (Fig. 2D,H,L-Q), interpretation is complicated by the evidence for dysfunction. Nonetheless, these experiments demonstrate a correlation between disrupted antigen-activation and disrupted changes in membrane topography.

#### Membrane topography correlates localization of ordered-lipid nanodomains and signaling proteins

Previous studies established that antigen-clustering of FcεRI in RBL mast cells causes proximal stabilization of ordered-lipid nanodomains, which facilitate functional coupling with Lyn kinase to initiate transmembrane signaling (22, 27, 47). A similar mechanism has been described for immune receptors in B cells and T cells (21, 24, 25). Localization of ordered-lipid nanodomains with microvilli was recently indicated for T cells (12). However, how these lipid domains and microvilli coordinate to localize signaling proteins and modulate their distribution after antigen activation for any immune cell has not been described. Several genetically encoded constructs have been developed to serve as lipid probes, including fluorescent proteins fused with fatty acylated peptides that anchor these probes to the inner leaflet of the plasma membrane (22, 27, 55). The PM lipid probe incorporates a short N-terminal segment of Lyn, including its palmitoylation and myristoylation sites that cause preferential localization to ordered-lipid regions. The GG lipid probe, commonly used for partitioning preferentially into disordered-lipid regions, contains the membrane anchorage motif like that of K-Ras consisting of a geranylgeranyl modification and a juxtamembrane polybasic sequence.

We employed the PM lipid probe to identify the distribution of ordered-lipid nanodomains with respect to the membrane topography of RBL mast cells. We prepared a 3xHA-tagged construct of PM, which could be labeled with fluorescent anti-HA antibodies. After transfection of the 3HA-PM plasmid, cells were stained with FM143fx, fixed, permeabilized, and labeled with anti-HA conjugated to Alexa 647. Results after MVC imaging are shown in Fig. 3 and SI Fig. S3. We observed that membrane topography of RBL mast cell does not change significantly upon transfection of PM and GG probes as can be quantified by our analysis of total microvillar area relative to the total cell area (SI Fig. S3F). We found that 93±1% and 90±2% of PM probes localize to the microvilli in the 0 nm and 400 nm slice, respectively (Fig. 3A,E; SI Fig. S3A,E). PM probes occupy 81±4% of the microvilli (Fig. 3F). Moreover, the δCount/δArea *vs* R plot for PM probes decreases sharply as the distance from the central microvillar region increases (Fig. 3G). This strong localization of PM probes indicates that the ordered-lipid nanodomains are found predominately in microvilli of suspended RBL mast cells.

**Figure 3:**
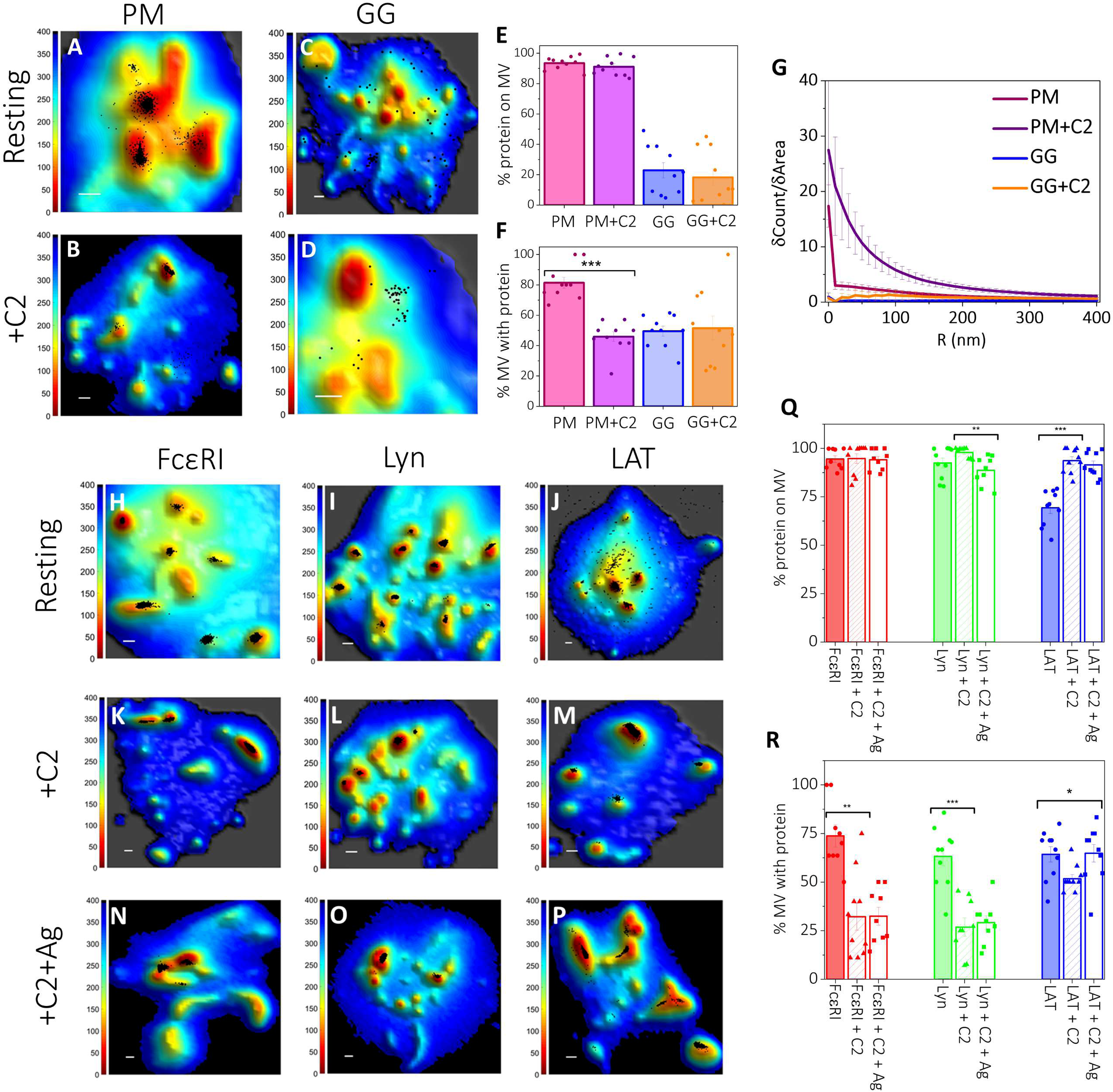
Synergy between Membrane Topography and Membrane Domains Controls the Localization of Signaling Proteins in Resting and Antigen-Activated States of RBL Mast Cells: A-D. Localization maps of probes preferring either ordered lipid (PM-3HA; A-B) or disordered lipid (3HA-GG; C-D) regions with respect to 3D membrane topography in resting untreated (A, C), and C2-ceramide treated (B, D) cells. Positions of probes obtained from SLN (black dots) in the 0 nm imaging slice are superimposed on membrane topography maps obtained from VA-TIRFM. The color bars represent the z distance from the glass in nanometers (nm). Scale bars (x,y), 0.5 μm. E-G. Quantified distributions of lipid probes on the RBL cell surface in resting untreated and C2-ceramide treated cells. The error bar represents the standard error of the mean. E. Percentage of specified lipid probes on microvillar (MV) regions of the membrane in resting untreated and C2-ceramide treated cells at 0 nm imaging plane. The values for individual cells are shown as dots in the plot. F. Fraction of microvilli (MV) occupied by the respective lipid probes in resting untreated and C2-ceramide treated cells. The values for individual cells are shown as dots in the plot. G. Probe density (δCount/δArea) as a function of radial distance (R) from the central microvilli region in the 0 nm imaging slice for resting untreated and C2-ceramide treated cells. H-P. Localization maps of specified proteins with respect to 3D membrane topography in resting untreated (H-J), C2-ceramide treated sensitized cells (K-M) and after 15 min antigen-activation in continued presence of C2-ceramide (N-P). Positions of proteins obtained from SLN (black dots) in the 0 nm imaging plane are superimposed on membrane topography maps obtained from VA-TIRFM. The color bars represent the z distance from the glass in nanometers (nm). Scale bars (x,y), 0.5 μm. Q-R. Quantified distributions of proteins in untreated, C2-ceramide treated resting cells, and after 15 min antigen-activation in continued presence of C2-ceramide. The error bar represents the standard error of the mean. Q. Percentage of specified proteins on membrane microvilli (MV) in untreated and C2-ceramide treated resting cells, and after 15 min antigen-activation in continued presence of C2-ceramide in the 0 nm imaging slice. The values for individual cells are shown as dots in the plot. R. Fraction of microvilli (MV) occupied by the specified proteins in untreated and C2-ceramide treated resting cells, and after 15 min antigen-activation in continued presence of C2-ceramide. The values for individual cells are shown as dots in the plot. P-values for Q and R are given in SI Table S2.

Using a 3xHA derivative of the disorder-preferring GG lipid probe, we found that the distribution of GG-3HA differs markedly from that of 3HA-PM. Only 23±5% and 18±5% of GG probes localize to the microvilli of suspended cells in 0nm and 400 nm slices, respectively (Fig. 3C,E; Fig. S3C,E). The fraction of microvilli occupied by the GG probes is 49±3% compared to 81±4% for PM probes (Fig. 3F). These partitioning values suggest that disordered-lipid regions preferentially occupy the flatter membranes of the cell body. The flat δCount/δArea *vs* R plot for GG probes also indicates that GG is spread more evenly across the cell membrane (Fig. 3G).

For more information about probe localization we employed N-acetyl-D-sphingosine (C2-ceramide) which like some other short-chain ceramides have been shown to disrupt ordered-lipid domains in model (56, 57) and cell (58–60) systems. For example, we showed that C2-ceramide causes concentration-dependent decreases in the lipid ordered regions of giant plasma membrane vesicles isolated from RBL cells and reduces the level of fluorescence resonance energy transfer between ordered-lipid-associating probes on intact RBL cells. The biologically inactive C2-ceramide analog, C2-dihydroceramide, does not have these effects (59). We further related disruption of IgE-FcεRI signaling mediated by ordered-lipids to C2-ceramide disruption of these nanodomains (59, 60). General effects of short chain ceramides on cellular physiology have been reviewed (61), and induction of apoptosis of cultured bone marrow-derived mast cells with 30 μM C2-ceramide has been reported but only after 6-hour treatment (61). Our treatment of RBL cells with 32 μM C2-ceramide was limited to a short time (∼10 min), which we have shown to be effective for disrupting lipid ordering in the plasma membrane (59, 60) but not likely to affect activities not related to ordered-lipid domains and not affecting cell survival (61).

MVC analysis showed that our C2 ceramide treatment of resting RBL cells does not cause a significant change in percentage microvillar area (13±1%, P = 0.39) indicating no substantial change in the microvilli or their density (SI Fig. S3N). We found the percentage of PM probes localizing to microvilli of C2-ceramide treated cells is also like that of untreated cells (∼90%; Fig. 3B,E; SI Fig. S3B,E). However, the fraction of microvilli occupied by PM probes decreases by 45% (from 81±4% to 46±3%) for C2-ceramide treated cells compared to untreated cells (Fig. 3F). In other words, C2-ceramide apparently causes PM probes to become more concentrated on fewer microvilli. Notably, the distribution of the GG probe does not change significantly upon C2-ceramide treatment of resting cells (Fig. 3D-G).

#### Perturbation of ordered-lipid nanodomains affects topographical localization of signaling proteins and degranulation

As described above, disruption of the ordered-lipid nanodomains has proven useful for investigating their functional relevance (58–60). We treated RBL cells with C2-ceramide to determine effects on the topographical localization of FcεRI, Lyn, and LAT, before and after activation with antigen. We found that, compared to untreated cells (Fig. 2I-K), antigen addition to C2-ceramide-treated cells for 15 min does not cause a dramatic topographical change, MVC imaging showing only limited merger of microvilli (Fig. 3N-P). Antigen activation of C2-ceramide-treated cells appears to cause a modest increase in the percentage microvillar area, although the activated value is not statistically different from that of untreated resting cells (SI Fig. S3N). These observations suggest that C2-ceramide-treated RBL cells do not activate normally after engagement with antigen, which is consistent with our previous findings that C2-ceramide inhibits antigen-stimulated signaling events including Ca^2+^ mobilization and phospholipase D activity (59, 60). To evaluate further, we measured antigen-stimulated degranulation of cells with and without C2-ceramide treatment. We found C2-ceramide essentially prevents stimulated degranulation (SI Fig. S3O), consistent with the disruption in treated cells of ordered-lipid nanodomains necessary to initiate signaling. We also tested localization of signaling proteins as follows.

#### FcεRI

For resting cells treated with C2-ceramide, FcεRI still localizes preferentially to the microvilli (95±2% and 95±2% on the 0 and 400 nm slices, respectively) like untreated cells (Fig. 3K,Q; SI Fig. S3G,M). However, the fraction of microvilli occupied by FcεRI decreases by 60% from 74±6% to 32±7%, after C2 ceramide-treatment (Fig. 3K,R), consistent with the substantial functional disruption by C2-ceramide. We found that 15 min antigen-activation of C2-ceramide-treated cells does not significantly change the percentage of FcεRI on the reduced fraction of occupied microvilli (94±2% and 89±3% on 0 and 400 nm slices, respectively (Fig. 3N,Q; SI Fig. S3J,M). The percentage of microvilli occupied by FcεRI after activation does not change from resting, C2-ceramide treated cells (Fig. 3K,N,R).

#### Lyn

The distributional responses of Lyn to C2-ceramide treatment are very similar to those of FcεRI. For resting cells, there is very little effect on Lyn’s predominant localization to microvilli (98±1% and 94±2% in the 0 nm and 400 nm slices, respectively, (Fig. 3L,Q; SI Fig. S3H,M).

Further, the percentage of microvilli occupied by Lyn is markedly reduced to less than half that of untreated cells (63±5% to 27±5%, Fig. 3R). Notably, the C2-ceramide-induced decrease in the fraction of microvilli occupied by Lyn parallels the decrease in the fraction of microvilli occupied by the ordered-lipid preferring PM probes in resting cells (Fig. 3F). Lyn still localizes primarily to microvilli after 15 min antigen-activation (89±3% and 87±3% in 0 nm and 400 nm slices, respectively (Fig. 3O,Q; SI Fig. S3K,M), and the fraction of microvilli occupied by Lyn kinase (29±4%, Fig. 3R) also remains the same as the C2-ceramide-treated, resting cells.

#### LAT

In contrast to FcεRI and Lyn, LAT shows ∼1.5-fold increased localization on the microvilli (P = 2 x 10^-6^) in C2-ceramide-treated, resting cells compared to untreated cells: 69±3% to 94±2% (Fig. 3 J,M,Q) and 68±3% to 91±3% (SI Fig. S3I,M) in 0 nm and 400 nm slices, respectively. The percentage of microvilli occupied by LAT molecules decreases from 64±4% to 52±2% in C2-ceramide-treated resting cells compared to untreated resting cells ((P = 0.01) Fig. 3R), a considerably smaller reduction compared to FcεRI and Lyn. 15 min after antigen addition to C2-ceramide-treated cells, LAT continues to localize predominantly to microvilli, 91±2% Fig. 3P,Q) and 91±2% (SI Fig. S3L,M) in 0 nm and 400 nm slices, respectively. The fraction of microvilli occupied by LAT appears to increase slightly after antigen-activation of C2-ceramide-treated cells (to 65±5%, Fig. 3R). This variable behavior of LAT after C2-ceramide treatment is not readily interpreted but may reflect previous observations that the order-preference of this scaffold protein is related to its palmitoylation state (49), which is unknown under these conditions.

#### Lyn and FcεRI appear to localize to mutually exclusive microvilli after C2-ceramide treatment

Our observation that both FcεRI and Lyn are restricted to roughly half the number of microvilli after C2-ceramide treatment raises the question whether both localize to the same microvillus or to different microvilli. If they localize to the same microvillus, then we might expect increased interactions and thereby enhanced cell activation, which we do not observe. To investigate we performed co-localization probability (CP) analysis (4, 5, 11) for FcεRI and Lyn in resting RBL cells that were treated or not with C2-ceramide. For that purpose, we labeled FcεRI with Alexa-647 tagged IgE and Lyn with Alexa-488 tagged anti-Lyn antibody. After our usual procedure to fix and label the suspended cells, we performed sequential dual-color SLN imaging to assess the CP values for these molecules in pairs on cells with nanometer resolution (Fig. 4). The distance-dependent CP is the probability that a probe of one type will have at least one partner of the other type within a specified interaction distance. The CP for probes *a* and *b* within distance R is given by CP = N*ab*(R)/N*a*, where N*a* is the total number of detected points of probe *a*, and N*ab*(R) is the number of points of probe *a* that have at least one partner of molecule *b* within R. We found that for resting cells at least ∼50% of FcεRI have one Lyn within 100 nm, i.e., within the same microvillus (Fig. 4A-C,G). The CP value increased to 80% at 400 nm. However, the fraction of IgE-FcεRI that have one Lyn within 20 nm distance is below 5% for resting cells (Fig. 4G). This indicates that although nearly half of the IgE-FcεRI localize with Lyn kinase molecules within a single microvillus, they are in separate assemblies. This might play a crucial role in controlling the auto-activation of mast cells in the resting state. Notably, in C2-ceramide treated cells, only ∼ 5% of FcεRI receptors have one Lyn kinase molecule within 100 nm distance, rising to 40% at 400 nm. (Fig. 4D-G).

**Figure 4:**
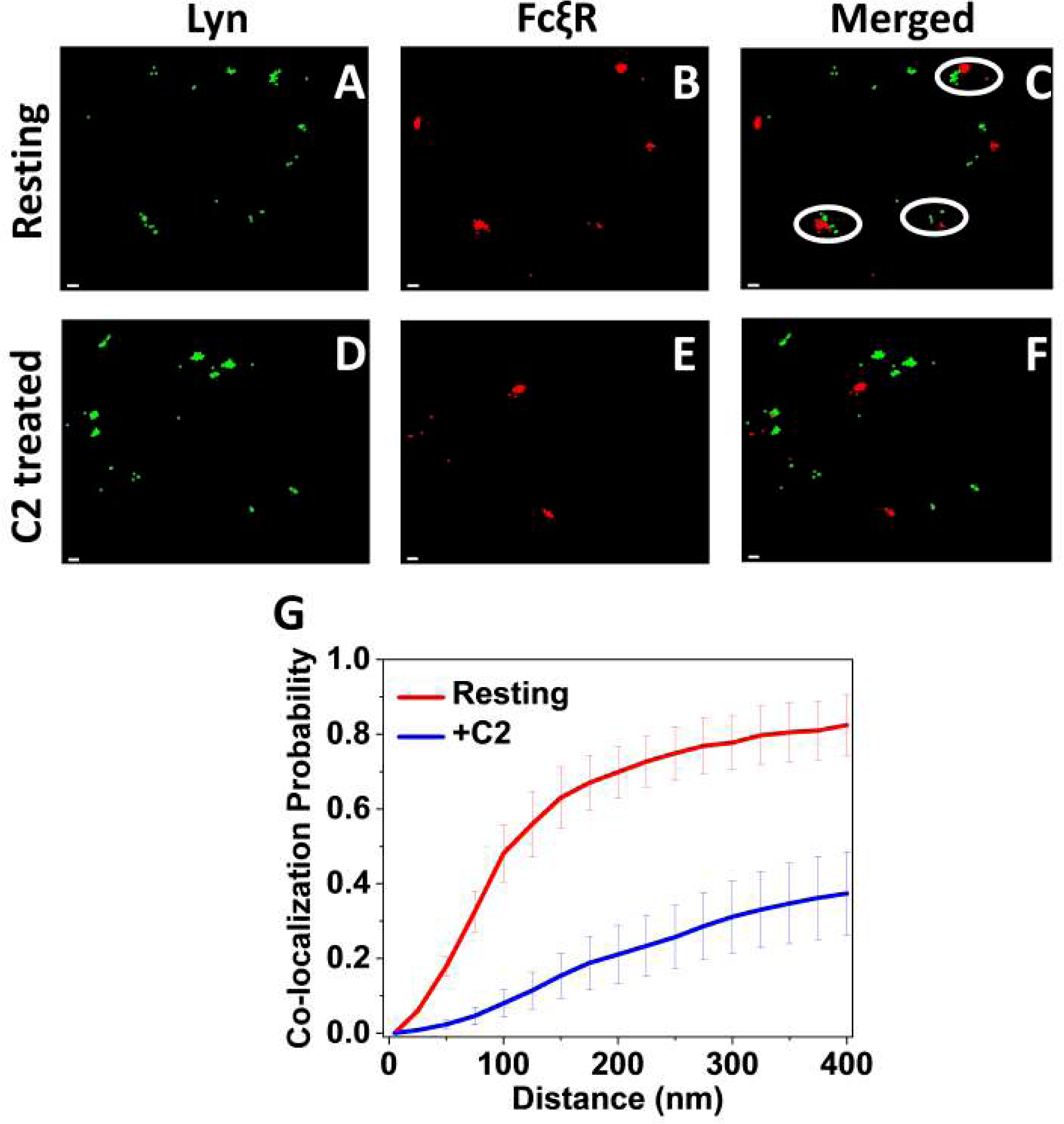
Colocalization Probability (CP) Analysis Reveals Close but Separate Concentrations of Lyn and FcξRI within Individual Microvilli of Resting RBL Mast Cells that Appear to Relocate to Mutually Exclusive Microvilli upon C2-Ceramide Treatment: A. SLN image of RBL cell labeled with Alexa Fluor-488-conjugated anti-Lyn antibodies (green), which stain Lyn. B. SLN image of the same cell labeled with Alexa Fluor 647-IgE (red), which stains FcεRI. C. Overlay of A and B. The white circle indicates where concentrations of Lyn and FcεRI are very close but not merged. D. SLN image of C2-ceramide-treated RBL cell with Lyn labeled as in A (green). E. SLN image of the same cell with FcεRI labeled as in B (red). F. Overlay of D and E. Scale bar 50 nm. G. CP as a function of distance for the pairs of Lyn and FcεRI in resting untreated and C2-ceramide treated RBL cells, obtained from analysis of five double-labeled cells for each condition. Error bars represent standard error of the mean.

Although we cannot rule out localization to extreme ends of extended microvilli, our results indicate that C2-ceramide treatment causes the majority of IgE-FcεRI and Lyn kinase molecules localize to mutually exclusive microvilli. Results from all our MVC, SLN, and CP experiments are summarized in the schematic of SI Fig. S4.

## Discussion

### FcεRI concentrates on RBL mast cell microvilli which merge to form ruffles upon antigen activation

Like B cells, T cells, and lymphocytes more generally (35), the plasma membrane of resting mast cells is dominated by dynamic, actin-mediated, finger-like protrusions known as microvilli (15–19). Previous studies demonstrated concentration of immune receptors in microvilli of B cells and T cells could be directly related to antigen sensing (2, 3, 5, 6, 8, 9, 12). The present study investigated whether the same is true in mast cells which play multiple roles in immune responses, including inflammation and allergies. We examined RBL mast cells, which have proved to be an experimentally robust model for examining the mechanisms of mast cells more generally (36). The topography of these RBL cells also resembles primary mast cells (15–18). For example, Hide et al used SEM to image the highly microvillar surface of resting peritoneal mast cells from Sprague-Dawley rats, as well as the ruffles that form upon 48/80 stimulation leading to degranulation (15). Using both SEM and transmission electron microscopy, Espinosa-Riquer et al. showed the microvillar membrane topography of resting bone marrow-derived mast cells and further showed the surface morphology changing to ruffles upon stimulation with antigen (18). In confocal images of resting BMMCs Matsuda et al. (62) showed fluorescent puncta corresponding to FcεRI, a pattern demonstrated in other cell types to signify microvilli-selective localization (63).

For RBL mast cells, the presence of microvilli that transform to membrane ruffles after antigen-activation has been observed with SEM and fluorescence microscopy (19, 39), but participation of membrane topography in antigen sensing and FcεRI-mediated signaling in any type of mast cell has been largely unexamined. Recent studies using confocal microscopy, MVC and lattice light-sheet microscopy (LLSM) highlighted the role of microvilli to present immune receptors for optimal antigen surveillance and to facilitate stimulated signaling pathways in T-cells and B-cells (2, 3, 5, 6, 8, 9, 12). Our present study extends this investigation by building on extensive studies that have been carried out on signaling in RBL mast cells initiated by FcεRI. We now advance this characterization as enabled by MVC, which with VA-TIRFM can nanoscopically image the three-dimensional structure of microvilli and with SLN can localize selected probes with respect to these topographically defined microvilli. Thus, we could correlate changes in plasma membrane topography of cells with distributions of signaling proteins as well as participation of ordered-lipid (Lo-like) nanodomains (colloquially known as “rafts”), which coordinately regulate and target antigen-induced signal initiation.

Our present study examined suspended RBL mast cells and the possibility that their microvilli pre-organize FcεRI and other membrane components to facilitate transmembrane signaling upon addition of antigen. Previous studies using this MVC approach with human effector and Jurkat T cells showed that TCR, co-receptor CD4, tyrosine kinase Lck, and adaptor LAT are substantially enriched in the microvilli on resting cells (2, 5), providing strong evidence that this level of localization yields a signaling hub that is key to T cell capacity to surveil juxtaposed antigen presenting cells (34, 64). Confirming a tight spatial correlation between microvilli and P-ERM (65, 66), Ghosh et al showed that these actin attachment proteins are necessary for localization of TCR to microvilli on resting cells (5). We found that the microvillar structure as defined by MVC is similar in the resting state for both T cells and RBL mast cells, and moreover that FcεRI and signaling partners concentrate in these structures. Like T cells and B cells, this level of organization may serve to enhance the capacity of resting mast cells to respond quickly and efficiently to environmental antigens.

Ghosh et al (5) further showed with MVC that addition of anti-CD3 to activate TCR on Jurkat cells as they interacted with an ICAM coated surface caused cells to spread quickly on those surfaces within 3 min (5). In contrast, our MVC analysis revealed that RBL cell microvilli do not quickly collapse on an antigen-presenting surface or in the presence of soluble antigen but rather transform within a few minutes to ridge-like ruffles. In comparison, Cai et al used diffraction-limited LLSM images of T cells to monitor the dynamics of microvilli, showing TCR clusters on the microvillar structures of resting cells and on actin-independent membrane protrusions within synapses (3). Park et al used high resolution confocal microscopy to demonstrate TCR on the tips of microvilli that were shed distally when these T cells formed synapses with activating surfaces (12). LLSM examination of Ramos B cells revealed dynamic microvilli interconnected by ridge-like structures (8). IgM-BCRs localize to the tops of these microvilli, evidently by an actin-driven mechanism, and antigen-induced clustering causes their immobilization on the ridges. Thus, antigen activation of RBL mast cells induces microvillar dynamics that differ significantly in comparison to those occurring with their immune cell counterparts, T cells or B cells.

### FcεRI and Lyn kinase localize to microvilli in separate protein assemblies that come together with LAT upon antigen activation

Using MVC to examine the distribution of endogenous signaling partners with respect to the 3D membrane topography of RBL mast cells, we found that IgE-FcεRI and Lyn, localize almost entirely in the microvilli on resting cells (Fig. 2). Our TIRF images of fluorescently labeled, suspended RBL cells showed consistent results (SI Fig. S2). This microvillar-specific localization resembles that of T cells for which TCR (analogous to FcεRI), Lck (analogous to Lyn), CD2, and CD4 are >80% localized to microvilli (5) . We found that antigen engagement causes the microvilli structures to merge into ruffles and that both FcεRI and Lyn continue to localize almost exclusively on these protrusions (Fig. 2 and SI Fig. S2). Our results are summarized in the schematic of SI Fig. S4.

Microvillar localization of both FcεRI and Lyn on resting mast cells raises the question how auto-activation in the absence of antigen is avoided. Our δCount/δArea *vs* R plot from MVC analysis shows slightly shifted peaks (Fig. 2) indicating that although occupying the same microvillus, FcεRI and Lyn maintain an inherent separation of a few tens of nanometers in the resting state. Further assessment with co-localization probability (CP) analysis clearly demonstrates that, although at least 80% of FcεRI have at least one Lyn partner within 400 nm (i.e. within a single microvillus) in the resting state, the percentage of FcεRI having a Lyn partner within very close proximity, i.e. within 20 nm, is very small (<5%) (Fig. 4). Thus, despite co-localization on the same microvillus in resting cells, this distinctive separation of FcεRI and Lyn is likely important for preventing spontaneous activation of mast cells. Upon antigen-crosslinking of FcεRI, these receptors are known to co-cluster with Lyn (22, 27, 30, 67). Consistent with those established results, MVC analysis shows overlapping δCount/δArea curves, indicating their merger (Fig. 2).

As for T cells, LAT in activated RBL mast cells serves as a scaffold for multiple signaling components, including Grb2, Gads, and SLP76 which connect to phosphorylated LAT via their SH2 motifs and connect to phospholipase-Cγ and other signaling enzymes (44, 68, 69). We determined for resting RBL cells that the larger fraction of LAT proteins localize to microvilli, but a significant fraction (∼30%) is found on the flatter cell body surface (Fig. 2). This distribution of LAT proteins in resting RBL mast cells is very similar to that in resting T-cells (5) with possible contributions from LAT within intracellular vesicles that exchange with the plasma membrane (70). By a minute after antigen-activation of RBL cells, almost all the LAT molecules localize to the merging microvillar structures (>90%), and the population in non-microvillar regions drastically decreases.

Correspondingly, the steepness of δCount/δArea vs R plot increases dramatically (Fig. 2). Remarkably, the shapes of δCount/δArea plots for IgE-FcεRI, Lyn, and LAT are very similar, indicating that these three proteins co-assemble upon activation. These results are consistent with our previous backscatter detection of immuno-gold particles in SEM images of adherent RBL cells. These dorsal surface images showed the separation of FcεRI, Lyn, and LAT in the resting state of mast cells and their co-clustering after antigen-crosslinking of FcεRI (67)

Because exceeding the phosphorylation threshold for antigen-crosslinked FcεRI to initiate cytoplasmic signaling involves the exclusion of transmembrane phosphatases from these clusters (27, 71), we attempted to investigate their localization in resting and activated cells. For this purpose, we employed genetically encoded PTPα-HA as a good proxy for these phosphatases in RBL mast cells (46), analogous to CD45 which operates in T cells. As monitored with fluorescent anti-HA antibodies, we found that PTPα-HA transiently transfected into resting RBL cells distributed randomly on the membrane surface without apparent selectivity for membrane topography (Fig. 2). We further observed that those cells with a high level of HA-PTPα expression do not undergo the signature membrane topographical changes to ruffles when exposed to antigen and that HA-PTPα transfected cells exhibit significantly reduced degranulation (SI Fig. S2). Although our experimental results with HA-PTPα may not represent the topographical distribution of endogenous transmembrane phosphatases, they are consistent with the requirement of FcεRI-mediated activation for changes in membrane topography that accompany downstream signaling in suspended cells.

### Plasma membrane topography coordinates with ordered-lipid nanodomains to regulate localization in resting and activated cells

Correlation of microvilli and ordered-lipid nanodomains was recently indicated for T cells by Park et al who found that microvilli bearing TCR are also decorated by added cholera toxin B, which binds to order-preferring ganglioside GM1(12). For a more thorough investigation of nanodomains in RBL mast cells, we employed genetically encoded probes that have been established to partition preferentially into ordered-lipid (palmitate/myristoylate: PM probe) or disordered-lipid (geranylgeranyl: GG probe) regions of the plasma membrane (55). With MVC analysis we found that 3HA-PM and GG-3HA localize differentially with respect to membrane topography in resting cells (Fig. 3). The ordered-lipid probe localizes predominantly with the microvilli, like Lyn and FcεRI. In contrast, the disordered-lipid probe prefers the flat, cell-body regions (Fig. 3 and as depicted in SI Fig. S4). Thus, these results demonstrate that ordered-lipid nanodomains tend to form on plasma membrane microvilli, whereas the membrane in the flat regions is more disordered in character. C2-ceramide treatment to disrupt ordered-lipid nanodomains (58–60) allowed us to evaluate possible coordination between these domains and actin-mediated membrane microvilli to control localization of FcεRI and Lyn. Although this perturbation does not significantly alter the microvillar-specific localization of FcεRI and Lyn, the fraction of microvilli occupied by each of those proteins decreases dramatically (by ∼50%) (Fig. 3). Remarkably, CP analysis indicates that those two proteins occupy mutually exclusive microvilli after C2-ceramide treatment (Fig. 4 and as depicted in SI Fig. S4). These results point to distinctive structural underpinnings for microvillus-selective localization of FcεRI and Lyn.

Lyn is known to partition into ordered-lipid regions as driven by dual palmitoylation and myristolyation of N-terminal amino acids (72), and 3HA-PM retains this segment of Lyn (55). Interestingly, the PM probe also reports that C2-ceramide mediated perturbation causes redistribution of the ordered-lipid domains to ∼50% of the microvilli, and this is likely to be the same set of microvilli as the 50% occupied by Lyn. Thus, for resting cells, the localization of Lyn and PM to microvilli is consistent with ordered-lipid nanodomains localized to these membranes. However, this is not a simple explanation for monomeric FcεRI, which exhibits little detectable preference for ordered-lipid regions before crosslinking by multivalent antigen leads to proximal stabilization of ordered-lipids (22, 47). This question is considered further in the next section.

### Organization of membrane proteins and lipids in microvilli may be mediated by the actin-cytoskeleton

It is known that parallel actin filaments create microvilli, and their stability is scaffolded by active (phosphorylated) ERM proteins (P-ERM), which are located almost exclusively with the microvillar structures (5, 66). Serving as a counterbalance is active cofilin, which severs the actin cytoskeleton upon dephosphorylation (33, 73, 74). We evaluated possible antigen-induced changes in these actin-attaching proteins that accompany changes in RBL mast cell signaling and membrane topography (SI Fig. S5 and S6). Our results show that by 1 min after antigen stimulation the phosphorylation of ERMs clearly outweighs dephosphorylation of cofilin (P-ERM/ERM > cofilin/P-cofilin), consistent with further enhancement of microvillar structures that are merging. By 15 min of stimulation the ratio of P-ERM/ERM compared to cofilin/P-cofilin decreases somewhat suggesting that the membrane ruffling occurring at this stage involves a balance of PERM/cofilin differing from that optimal for maintaining individual microvilli. C2-ceramide treatment prevents the increase of both the phosphorylated and total ERM after antigen exposure, consistent with preventing FcεRI-mediated cell activation by disrupting the coupling with Lyn. Our results with RBL cells can be compared to those for DT40 B cells which also showed that a consequence of antigenic stimulation is modulating the phosphorylation pattern of ERM proteins and cofilin.

However, different from the RBL cells, simultaneous dephosphorylation of ERM and cofilin in stimulated DT40 B cells leads to local depletion of actin, diminished microvillar structures, and enhanced BCR clustering (33).

We speculate that actin attachment proteins participate in the nearly exclusive localization of Lyn and FcεRI to microvilli in resting mast cells. We previously evaluated the segregation of ordered-lipid vs disordered-lipid regions of the plasma membrane as may be mediated by actin attachments (22). Our SLN measurements in those RBL cell studies are generally consistent with the findings of others that order-preferring attachment proteins can mediate actin contacts with order-preferring membrane proteins in lymphocyte signaling (75, 76). For example, Cbp/PAG is strongly order-preferring, a known interaction partner of Lyn, and coupled to the actin cytoskeleton via interactions with EBP-50 and ezrin (77). In this view, actin aligned in the microvilli might stabilize ordered-lipid nanodomains in the adjoining plasma membrane, as mediated in part by P-ERM and PAG-like associations. These nanodomains would also then preferentially include palmitoylated/myristoylated Lyn and other saturated fatty acid anchors (e.g., PM-3HA). That Lyn and FcεRI segregate to separate microvilli after C2-ceramide underscores differential actin association by FcεRI. Crosstalk between the FcεRI and ERM proteins has been observed experimentally in mast cells (78), but specific structural associations have not been identified.

### Microvilli facilitate antigen-sensitive transmembrane signaling in mast cells to cause functional morphological changes

Our nanoscopic MVC examination of RBL mast cells showed that, like T cells, the antigen receptor (FcεRI) concentrates on microvilli together with its key signaling partner (Lyn kinase), providing strong evidence that this assembly serves as an efficient sense-and-response mechanism for antigenic stimuli (5, 34, 64). Antigen activation of T cells, B cells, and mast cells is followed by topographical changes such as those occurring on antigen presenting cells or other activating surfaces. As shown for B cells (33) and RBL mast cells (present study), one consequence of cell activation is modulating the phosphorylation pattern of ERM proteins, which in turn modulates actin attachment in the microvilli (5). Current studies indicate differences among these three types of immune cells regarding the nature of the microvilli collapse (or distal molting) at an activating surface, although these apparent differences may be derived in part from the imaging approach employed (e.g., MVC vs LLSM vs confocal microscopy) (5, 8, 12). Our present MVC studies on RBL mast cells can be compared directly to those carried out on T cells (5), which showed rapid microvillar collapse on an activating surface (5). In contrast, mast cell microvilli merge into ruffles within a minute after engaging a surface-bound or soluble antigen (Fig 1 and 2).

Although MVC shows that flattening of RBL mast cells on surfaces is delayed compared to T cells, flattening and adherence do occur after about 30 min, as described previously (39, 44). The adherence process involves integrins, but FcεRI-mediated signaling does not occur in the absence of antigen (39, 44). Antigen activation causes the adherent cells to spread out and ruffles appear on the periphery (39, 44). Subject to further investigation is if and how the spatial relationships and structural associations of FcεRI, Lyn, LAT and ordered-lipid nanodomains change when microvilli flatten on the ventral surface of adherent cells. Overall, our MVC results are consistent with previous extensive SLN and ImFCS imaging carried out in TIRF mode on the ventral surface (22, 27) and SEM imaging on the dorsal surface (67), showing that antigen crosslinking of FcεRI stabilizes an ordered-lipid environment wherein coupling with Lyn and LAT occurs.

Although antigen-activated ruffling has long been observed in mast cells, its physiological role has not been clearly defined. Membrane ruffling in RBL mast cells can be separated from cell activation pathways necessary for degranulation (19, 79): It can be stimulated by phorbol esters, which mimic the PIP2 metabolite diacyl glycerol and act as an agonist for protein kinase C. Interestingly, PKC causes exposure of PIP2, implicated in antigen-stimulated phosphorylation of ERM proteins in B cells (33). However, phorbol ester-stimulated ruffling in is Ca^2+^ independent, whereas antigen-stimulated signaling leading to PKC activation and degranulation also requires Ca^2+^ mobilization (19, 79). Stimulated ruffling may be productively involved in mast cell motility. This possibility is consistent with our previous characterization of RBL mast cells moving up a chemical gradient of antigen (80). At low antigenic concentrations within the gradient, the motile cells move toward the source, but at a threshold concentration the cells adhere, spread, and begin to degranulate. Mast cells are known to migrate toward source of infection, making physiological sense that membrane dynamics facilitate motion until the stimulus becomes sufficiently strong to cause proximal degranulation to release of chemical mediators of inflammation.

## Materials and Methods

MVC methodology images and dual color super-resolution images were recorded using a homebuilt TIRF microscope equipped with an electron multiplying charge-coupled device (EMCCD) camera (5, 11, 27). The raw images were further processed using custom-made MATLAB codes (11) to determine the quantitative distribution of membrane protein as related to membrane topography and co-localization probability. The flow cytometry experiments were conducted on a Thermo Fisher Attune NxT flow cytometer. The SEM experiments were carried out on a Keck Leo 1550 SEM. Detailed descriptions of the source of all materials including chemicals and plasmids, instrumental setups, sample preparation, image processing, and quantitative analyses are provided in the SI Appendix.

### Data sharing plans

The code used in the analysis is available in a previous publication (13). Data and documentation are available upon reasonable request from the corresponding authors.

## Supporting information

Material and Methods, FACS experiments,Supporting figure 1-6, Supporting Table 1-4

## Acknowledgments

This work is supported by the National Institute of General Medical Sciences (NIGMS) Grant R01GM117552. SG acknowledges Start-up Research Grant (SRG/2023/000147), SERB-India for financial support during manuscript preparation after experimental work was completed. This work made use of the shared instrumentation facility at the Cornell Center for Materials Research and the BRC Flow Cytometry Facility (RRID:SCR_021740) at the Cornell Institute of Biotechnology. The content is solely the responsibility of the authors and does not necessarily represent the official views of NIGMS or NIH.

## References

1. R. C. Cail, D. G. Drubin, Membrane curvature as a signal to ensure robustness of diverse cellular processes. Trends Cell Biol (2022). 10.1016/j.tcb.2022.09.004.

2. Y. Jung, et al., Three-dimensional localization of T-cell receptors in relation to microvilli using a combination of superresolution microscopies. Proceedings of the National Academy of Sciences 113 (2016).

3. E. Cai, et al., Visualizing dynamic microvillar search and stabilization during ligand detection by T cells. Science 356 (2017).

4. S. Ghosh, et al., CCR7 signalosomes are preassembled on tips of lymphocyte microvilli in proximity to LFA-1. Biophys J 120, 4002–4012 (2021).

5. S. Ghosh, et al., ERM-Dependent Assembly of T Cell Receptor Signaling and Co-stimulatory Molecules on Microvilli prior to Activation. Cell Rep 30, 3434–3447.e6 (2020).

6. H.-R. Kim, et al., T cell microvilli constitute immunological synaptosomes that carry messages to antigen-presenting cells. Nat Commun 9, 3630 (2018).

7. M. Aramesh, et al., Nanoconfinement of microvilli alters gene expression and boosts T cell activation. Proceedings of the National Academy of Sciences 118 (2021).

8. D. Saltukoglu, et al., Plasma membrane topography governs the 3D dynamic localization of IgM B cell antigen receptor clusters. EMBO J 42 (2023).

9. Y. Jung, L. Wen, A. Altman, K. Ley, CD45 pre-exclusion from the tips of T cell microvilli prior to antigen recognition. Nat Commun 12, 3872 (2021).

10. Y. Ben Sahel, G. Dardikman-Yoffe, Y. C. Eldar, S. Gosh, G. Haran, Super-Resolved Imaging of Early-Stage Dynamics in the Immune Response in 2021 IEEE International Conference on Image Processing (ICIP), (IEEE, 2021), pp. 3468–3472.

11. S. Ghosh, A. Alcover, G. Haran, “Microvillar Cartography: A Super-Resolution Single-Molecule Imaging Method to Map the Positions of Membrane Proteins with Respect to Cellular Surface Topography” in The Immune Synapse: Methods and Protocols, C. T. Baldari, M. L. Dustin, Eds. (2023).

12. J.-S. Park, et al., Trogocytic molting of T cell microvilli upregulates T cell receptor surface expression and promotes clonal expansion. Nat Commun 14, 2980 (2023).

13. D. D. Metcalfe, D. Baram, Y. A. Mekori, Mast cells. Physiol Rev 77, 1033–1079 (1997).

14. H. Metzger, The Receptor with High Affinity for IgE. Immunol Rev 125, 37–48 (1992).

15. I. Hide, et al., Degranulation of individual mast cells in response to Ca2+ and guanine nucleotides: an all-or-none event. J Cell Biol 123, 585–593 (1993).

16. Y. Romo-Lozano, F. Hernández-Hernández, E. Salinas, Sporothrix schenckii yeasts induce ERK pathway activation and secretion of IL-6 and TNF-α in rat mast cells, but no degranulation. Med Mycol 52, 862–868 (2014).

17. H. Lin, et al., Cytotoxicity assessment of exfoliated MoS 2 using primary human mast cells and the progenitor cell-derived mast cell line LAD2. Nanoscale Adv 6, 2419–2430 (2024).

18. Z. P. Espinosa-Riquer, et al., Signal Transduction Pathways Activated by Innate Immunity in Mast Cells: Translating Sensing of Changes into Specific Responses. Cells 9, 2411 (2020).

19. J. R. Pfeiffer, J. C. Seagrave, B. H. Davis, G. G. Deanin, J. M. Oliver, Membrane and cytoskeletal changes associated with IgE-mediated serotonin release from rat basophilic leukemia cells. J Cell Biol 101, 2145–2155 (1985).

20. E. Sherman, et al., Functional nanoscale organization of signaling molecules downstream of the T cell antigen receptor. Immunity 35, 705–20 (2011).

21. M. B. Stone, S. A. Shelby, M. F. Núñez, K. Wisser, S. L. Veatch, Protein sorting by lipid phase-like domains supports emergent signaling function in B lymphocyte plasma membranes. Elife 6 (2017).

22. S. A. Shelby, S. L. Veatch, D. A. Holowka, B. A. Baird, Functional nanoscale coupling of Lyn kinase with IgE-FcεRI is restricted by the actin cytoskeleton in early antigen-stimulated signaling. Mol Biol Cell 27, 3645–3658 (2016).

23. I. Levental, S. L. Veatch, The Continuing Mystery of Lipid Rafts. J Mol Biol 428, 4749–4764 (2016).

24. T. Harder, Lipid raft domains and protein networks in T-cell receptor signal transduction. Curr Opin Immunol 16, 353–359 (2004).

25. J. Dinic, A. Riehl, J. Adler, I. Parmryd, The T cell receptor resides in ordered plasma membrane nanodomains that aggregate upon patching of the receptor. Sci Rep 5, 10082 (2015).

26. N. Bag, E. London, D. A. Holowka, B. A. Baird, Transbilayer Coupling of Lipids in Cells Investigated by Imaging Fluorescence Correlation Spectroscopy. J Phys Chem B 126, 2325– 2336 (2022).

27. N. Bag, et al., Lipid-based and protein-based interactions synergize transmembrane signaling stimulated by antigen clustering of IgE receptors. Proceedings of the National Academy of Sciences 118 (2021).

28. M. Rouches, S. L. Veatch, B. B. Machta, Surface densities prewet a near-critical membrane. Proceedings of the National Academy of Sciences 118 (2021).

29. H.-Y. Wang, et al., Coupling of protein condensates to ordered lipid domains determines functional membrane organization. Sci Adv 9 (2023).

30. S. A. Shelby, D. Holowka, B. Baird, S. L. Veatch, Distinct stages of stimulated FcεRI receptor clustering and immobilization are identified through superresolution imaging. Biophys J 105, 2343–54 (2013).

31. D. C. Seldin, et al., Homology of the rat basophilic leukemia cell and the rat mucosal mast cell. Proceedings of the National Academy of Sciences 82, 3871–3875 (1985).

32. S. Faure, et al., ERM proteins regulate cytoskeleton relaxation promoting T cell–APC conjugation. Nat Immunol 5, 272–279 (2004).

33. A. Droubi, et al., The inositol 5-phosphatase INPP5B regulates B cell receptor clustering and signaling. Journal of Cell Biology 221 (2022).

34. R. Orbach, X. Su, Surfing on Membrane Waves: Microvilli, Curved Membranes, and Immune Signaling. Front Immunol 11 (2020).

35. S. Majstoravich, et al., Lymphocyte microvilli are dynamic, actin-dependent structures that do not require Wiskott-Aldrich syndrome protein (WASp) for their morphology. Blood 104, 1396– 403 (2004).

36. F. H. Falcone, D. Wan, N. Barwary, R. Sagi-Eisenberg, RBL cells as models for in vitro studies of mast cells and basophils. Immunol Rev 282, 47–57 (2018).

37. M. A. Govendir, et al., T cell cytoskeletal forces shape synapse topography for targeted lysis via membrane curvature bias of perforin. Dev Cell 57, 2237–2247.e8 (2022).

38. J. M. Oliver, J. Seagrave, R. F. Stump, J. R. Pfeiffer, G. G. Deanin, Signal transduction and cellular response in RBL-2H3 mast cells. Prog Allergy 42, 185–245 (1988).

39. R. N. Orth, M. Wu, D. A. Holowka, H. G. Craighead, B. A. Baird, Mast Cell Activation on Patterned Lipid Bilayers of Subcellular Dimensions. Langmuir 19, 1599–1605 (2003).

40. R. Rea, et al., Streamlined synaptic vesicle cycle in cone photoreceptor terminals. Neuron 41, 755–66 (2004).

41. M. D. Sharp, K. Pogliano, An in vivo membrane fusion assay implicates SpoIIIE in the final stages of engulfment during Bacillus subtilis sporulation. Proc Natl Acad Sci U S A 96, 14553–8 (1999).

42. Y. Fu, et al., Axial superresolution via multiangle TIRF microscopy with sequential imaging and photobleaching. Proceedings of the National Academy of Sciences 113, 4368–4373 (2016).

43. P. Sundd, et al., Quantitative dynamic footprinting microscopy reveals mechanisms of neutrophil rolling. Nat Methods 7, 821–4 (2010).

44. D. L. Wakefield, D. Holowka, B. Baird, The FcεRI signaling cascade and integrin trafficking converge at patterned ligand surfaces. Mol Biol Cell 28, 3383–3396 (2017).

45. B. Baird, D. Sajewski, S. Mazlin, A microtiter plate assay using cellulose acetate filters for measuring cellular [3H]serotonin release. J Immunol Methods 64, 365–375 (1983).

46. R. M. Young, X. Zheng, D. Holowka, B. Baird, Reconstitution of regulated phosphorylation of FcepsilonRI by a lipid raft-excluded protein-tyrosine phosphatase. J Biol Chem 280, 1230–5 (2005).

47. K. A. Field, D. Holowka, B. Baird, Fc epsilon RI-mediated recruitment of p53/56lyn to detergent-resistant membrane domains accompanies cellular signaling. Proc Natl Acad Sci U S A 92, 9201–5 (1995).

48. A. M. Davey, R. P. Walvick, Y. Liu, A. A. Heikal, E. D. Sheets, Membrane order and molecular dynamics associated with IgE receptor cross-linking in mast cells. Biophys J 92, 343–55 (2007).

49. I. Levental, D. Lingwood, M. Grzybek, Ü. Coskun, K. Simons, Palmitoylation regulates raft affinity for the majority of integral raft proteins. Proceedings of the National Academy of Sciences 107, 22050–22054 (2010).

50. S. Saitoh, et al., LAT is essential for Fc(epsilon)RI-mediated mast cell activation. Immunity 12, 525–35 (2000).

51. A. Geldman, C. J. Pallen, Protein tyrosine phosphatases in mast cell signaling. Methods Mol Biol 1220, 269–86 (2015).

52. R. A. Fernandes, et al., A cell topography-based mechanism for ligand discrimination by the T cell receptor. Proceedings of the National Academy of Sciences 116, 14002–14010 (2019).

53. S. A. Berger, T. W. Mak, C. J. Paige, Leukocyte common antigen (CD45) is required for immunoglobulin E-mediated degranulation of mast cells. J Exp Med 180, 471–6 (1994).

54. K. M. Chisholm, et al., Mast cells in systemic mastocytosis have distinctly brighter CD45 expression by flow cytometry. Am J Clin Pathol 143, 527–34 (2015).

55. P. S. Pyenta, D. Holowka, B. Baird, Cross-correlation analysis of inner-leaflet-anchored green fluorescent protein co-redistributed with IgE receptors and outer leaflet lipid raft components. Biophys J 80, 2120–32 (2001).

56. S. Chiantia, N. Kahya, P. Schwille, Raft Domain Reorganization Driven by Short- and Long-Chain Ceramide: A Combined AFM and FCS Study. Langmuir 23, 7659–7665 (2007).

57. Megha, P. Sawatzki, T. Kolter, R. Bittman, E. London, Effect of ceramide N-acyl chain and polar headgroup structure on the properties of ordered lipid domains (lipid rafts). Biochimica et Biophysica Acta (BBA) - Biomembranes 1768, 2205–2212 (2007).

58. A. Pacheco, et al., C2-Phytoceramide Perturbs Lipid Rafts and Cell Integrity in Saccharomyces cerevisiae in a Sterol-Dependent Manner. PLoS One 8, e74240 (2013).

59. A. Gidwani, H. A. Brown, D. Holowka, B. Baird, Disruption of lipid order by short-chain ceramides correlates with inhibition of phospholipase D and downstream signaling by FcɛRI. J Cell Sci 116, 3177–3187 (2003).

60. D. Holowka, K. Thanapuasuwan, B. Baird, Short chain ceramides disrupt immunoreceptor signaling by inhibiting segregation of Lo from Ld Plasma membrane components. Biol Open 7 (2018).

61. R. Ghidoni, G. Sala, A. Giuliani, Use of sphingolipid analogs: benefits and risks1The ganglioside nomenclature is that of Svennerholm, L. (1969) J. Lipid Res. 5, 145–155.1. Biochimica et Biophysica Acta (BBA) - Molecular and Cell Biology of Lipids 1439, 17–39 (1999).

62. A. Matsuda, et al., High-affinity IgE receptor-β chain expression in human mast cells. J Immunol Methods 336, 229–234 (2008).

63. P. Colarusso, K. R. Spring, Reticulated Lipid Probe Fluorescence Reveals MDCK Cell Apical Membrane Topography. Biophys J 82, 752–761 (2002).

64. L. Balagopalan, K. Raychaudhuri, L. E. Samelson, Microclusters as T Cell Signaling Hubs: Structure, Kinetics, and Regulation. Front Cell Dev Biol 8, 608530 (2020).

65. S. Yonemura, S. Tsukita, S. Tsukita, Direct Involvement of Ezrin/Radixin/Moesin (ERM)- binding Membrane Proteins in the Organization of Microvilli in Collaboration with Activated ERM Proteins. J Cell Biol 145, 1497–1509 (1999).

66. A. Hanono, D. Garbett, D. Reczek, D. N. Chambers, A. Bretscher, EPI64 regulates microvillar subdomains and structure. J Cell Biol 175, 803–813 (2006).

67. S. L. Veatch, E. N. Chiang, P. Sengupta, D. A. Holowka, B. A. Baird, Quantitative nanoscale analysis of IgE-FcεRI clustering and coupling to early signaling proteins. J Phys Chem B 116, 6923–35 (2012).

68. L. Balagopalan, R. L. Kortum, N. P. Coussens, V. A. Barr, L. E. Samelson, The Linker for Activation of T Cells (LAT) Signaling Hub: From Signaling Complexes to Microclusters. Journal of Biological Chemistry 290, 26422–26429 (2015).

69. M. A. Silverman, J. Shoag, J. Wu, G. A. Koretzky, Disruption of SLP-76 Interaction with Gads Inhibits Dynamic Clustering of SLP-76 and FcεRI Signaling in Mast Cells. Mol Cell Biol 26, 1826–1838 (2006).

70. A. Alcover, B. Alarcón, V. Di Bartolo, Cell Biology of T Cell Receptor Expression and Regulation. Annu Rev Immunol 36, 103–125 (2018).

71. D. Holowka, B. Baird, Roles for lipid heterogeneity in immunoreceptor signaling. Biochim Biophys Acta 1861, 830–836 (2016).

72. M. Kovářová, et al., Structure-Function Analysis of Lyn Kinase Association with Lipid Rafts and Initiation of Early Signaling Events after Fcɛ Receptor I Aggregation. Mol Cell Biol 21, 8318–8328 (2001).

73. B. W. Bernstein, J. R. Bamburg, ADF/Cofilin: a functional node in cell biology. Trends Cell Biol 20, 187–195 (2010).

74. M. S. Wang, M. Huse, Phollow the phosphoinositol: Actin dynamics at the B cell immune synapse. Journal of Cell Biology 221 (2022).

75. B. P. Head, H. H. Patel, P. A. Insel, Interaction of membrane/lipid rafts with the cytoskeleton: Impact on signaling and function. Biochimica et Biophysica Acta (BBA) - Biomembranes 1838, 532–545 (2014).

76. A. Viola, N. Gupta, Tether and trap: regulation of membrane-raft dynamics by actin-binding proteins. Nat Rev Immunol 7, 889–896 (2007).

77. M. Hrdinka, V. Horejsi, PAG - a multipurpose transmembrane adaptor protein. Oncogene 33, 4881–4892 (2014).

78. I. Hálová, et al., Cross-talk between Tetraspanin CD9 and Transmembrane Adaptor Protein Non-T Cell Activation Linker (NTAL) in Mast Cell Activation and Chemotaxis. Journal of Biological Chemistry 288, 9801–9814 (2013).

79. J. M. Oliver, J. C. Seagrave, J. R. Pfeiffer, M. L. Feibig, G. G. Deanin, Surface functions during mitosis in rat basophilic leukemia cells. J Cell Biol 101, 2156–66 (1985).

80. J. Lee, S. L. Veatch, B. Baird, D. Holowka, Molecular mechanisms of spontaneous and directed mast cell motility. J Leukoc Biol 92, 1029–1041 (2012).

