## Supplementary material for "Synergy between Membrane Topography and Domains to Control Signaling Protein Localization in Mast Cells Facilitates their Activation": Material and Methods, FACS experiments,Supporting figure 1-6, Supporting Table 1-4

#### 1. MATERIALS AND METHODS

**1.1. Reagents:** Minimum essential medium (MEM), Opti-MEM, Trypsin-EDTA (0.01%) and gentamicin sulfate were obtained from Life Technologies (Carlsbad, CA). Fetal Bovine Serum (FBS) was purchased from Bio-Techne (Minneapolis, MN). The antigen, DNP-BSA, was prepared by conjugating DNP sulfonate (Sigma Aldrich) to bovine serum albumin (BSA) (1). Alexa Fluor 647 (AF647) NHS ester (Invitrogen) was employed to fluorescently label monoclonal anti-DNP (2,4-dinitrophenyl) immunoglobulin E (IgE) yielding AF647-IgE as described previously (2). FM143 and FM143fx dyes and Alexa-568 labeled goat anti-rabbit secondary antibody were from Invitrogen. Rabbit antibodies specific for Ezrin/Radixin/Moesin, Phospho-Ezrin (Thr567)/Radixin (Thr564)/Moesin (Thr558) (48G2), Phospho-Cofilin (Ser3) (77G2) and Cofilin (D3F9) were purchased from Cell Signaling Technology (CST). Antibodies specific for HA-probe (F-7) Alexa Fluor® 647, Lyn (H-6) Alexa Fluor® 647, Lyn (H-6) Alexa Fluor® 488 and LAT (11B.12) Alexa Fluor® 647 were purchased from Santa Cruz Biotechnology (SCBT). Paraformaldehyde (16%) and Glutaraldehyde (25%) were purchased from Electron Microscopy Sciences (EMS). Poly-D-Lysine solution (1.0 mg/mL), phorbol 12,13-dibutyrate, cysteamine, glucose oxidase, catalase (Sigma), and glucose were purchased from Sigma-Aldrich.

**1.2. Plasmids:** 3HA-PM and GG-3HA were created from previously prepared constructs EGFP-PM and GG-EGFP (3) by performing a double digesting at restriction sites AgeI/NotI and AgeI/BglII, respectively, cutting out the sequence coding for EGFP and replacing with previously annealed oligonucleotides coding for a triple HA tag (IDT DNA). The PTPα-HA plasmid was a gift from David Shalloway (Cornell University) (3).

**1.3. Cell culture, transfection, sensitization, stimulation, and C2-ceramide treatments:**

*Cell culture:* RBL-2H3 mast cells (4, 5) (for brevity, RBL cells) were cultured for adherent growth in medium (80% MEM, supplemented with 20% FBS and 10 mg/L gentamicin sulfate) in a 37°C and 5% (v/v) $CO_2$ incubator. Cells were harvested with Trypsin-EDTA followed by sedimentation and resuspension in the specified medium as described previously (3).

*Chemical transfection.* About 20,000 cells were seeded in 2 mL growth medium in a 35 mm Petri dish and cultured overnight. Then cells were transfected using FuGENE HD transfection kit (Promega) following the manufacturer protocol. Briefly, the plasmid DNA (0.5 – 1 µg) and FuGENE (3 µL FuGENE/µg DNA) were first mixed in 100 µL Opti-MEM medium and incubated at room temperature for 15 min. The adherent cells in each dish were washed, then covered with 1 mL Opti-MEM. The DNA/FuGENE complex was added dropwise across the dish area, followed by incubation for 1 hr at 37°C. Dishes were then incubated with pre-warmed phorbol 12,13-dibutyrate (1 mL, 0.1 µg/mL) for 3 hr at 37°C in 5% (v/v) $CO_2$ environment. Following this incubation, the Opti-MEM was discarded, 2 mL of growth medium was added to each dish and the transfected cells were returned to the incubator for 18 – 22 hr before further processing.

*Cell sensitization and stimulation.* Confluent RBL cells in 25 $cm^2$ flask were washed twice with fresh media and then returned to the incubator for sensitization with 0.5 µg/mL anti-DNP IgE (labeled or not with Alexa-647) over night. Cells were then harvested and, after washing, resuspended at $3x10^6$/mL in growth medium containing 1 µg/mL DNP-BSA (antigen). These samples, non-stimulated controls, and non-sensitized cells were incubated at 37°C for 1 min or 15 min with slow rotation. Cells were then immediately treated with fixatives as described below for antibody labeling.

*C2 ceramide treatments.* Harvested cells were resuspended in media at $3x10^6$/mL containing 32µM C2-ceramide and incubated under slow rotation at 37°C for 10 min. Treated cells were activated by addition of DNP-BSA to a final concentration of 1 µg/mL and incubated at 37°C for 15 min under continued rotation. After this incubation in medium with or without (control) DNP-BSA or medium, cells were immediately treated with fixatives as described below for antibody labeling.

**1.4. Degranulation Assays:** RBL cells, transfected with PTPα as described above or not transfected, were harvested, resuspended in buffered salt solution (BSS: 135 mM NaCl, 5 mM KCl, 1.8 mM $CaCl_2$, 1.0 mM $MgCl_2$, 5.6 mM glucose, 20 mM HEPES, and 1 mg/mL BSA at pH 7.4), sensitized with 4 µg anti-DNP IgE/1mL of cells ($1x10^6$ cells/mL), and replated in a 96 well plate. After 2 hr incubation and washing, the standard degranulation plate assay was carried out in BSS as described previously (6). Stimulation was initiated by addition of antigen (10ng/mL DNP-BSA). Controls were (0.1% Triton-X 100) or BSS only to determine total and spontaneous release, respectively. After incubating the plate for 1 hr at 37°C, degranulation was terminated by placing the cells for 10 min on ice. To quantify the amount of β-hexosaminidase released by the cells, an aliquot of the supernatant was evaluated using β-hexosaminidase-substrate (4-methylumbelliferyl-*N*-acetyl-β-d-glucosaminide) (Millipore Sigma) in a fluorormetric assay (6). Background fluorescence with substrate in buffer alone was subtracted from all readings.

To evaluate effects of C2-ceramide treatments, cultured RBL cells were sensitized overnight with IgE (0.5 µg/ml), harvested the next day and washed with BSS. Suspended cells were divided in equal amounts into microtubes (Corning Inc) preparing 3 samples each containing RBL cells in BSS with one of the following: with antigen (10ng/ml DNP-BSA) only, with C2-ceramide (32µM) only, with C2-ceramide and antigen, with 0.1% Triton-X 100, or BSS only. Tubes were incubated at 37°C for 1h on a rotating device, placed on ice for 10 min and centrifuged in the cold for 5 min. Then the β-hexosaminidase release was quantified as described above.

**1.5. Cellular labeling with antibodies:** For resting cell samples, harvested cells were resuspended at $3x10^6$/mL and incubated on ice for 10 min in a blocking solution of phosphate-buffered saline (PBS) containing 1% (wt/vol) BSA, 5 mM EDTA. After sedimentation, the cells were washed twice with 5 mM EDTA/PBS by sedimentation and resuspension at 4°C. Then the cells were resuspended in a standard fixation buffer of PBS containing 4% (wt/vol) paraformaldehyde, 0.2–0.5% glutaraldehyde, 2% sucrose, 10 mM EGTA, and 1 mM EDTA. After incubation for 2 h on ice, the cells were sedimented and washed twice with PBS. For cells that were treated with C2-ceramide and/or activated with antigen DNP-BSA, the cell suspension was mixed with an equal volume of PBS fixation buffer, except containing 8% instead of 4% (wt/vol) paraformaldehyde (such that the effective concentration of paraformaldehyde after mixing is 4%). Cells were then immediately sedimented at 4°C and resuspended in the standard fixation buffer. After incubation for 2 h on ice, the cells were sedimented and washed twice with PBS as for the other samples.

We used FM143fx membrane dye (5 µg/mL) to homogeneously stain the plasma membrane of fixed cells by incubation for 30 min on ice as previously established (7–12). Cells were washed twice with the blocking buffer by sedimentation and resuspension at 4°C. Then the pellet was dissolved with the permeabilization buffer (0.1% TritonX-100, 2% BSA in PBS). An appropriate amount of fluorescently-labeled antibody specific for selected protein (~10 µg/mL) was added to the solution followed by incubation overnight at 4°C. Samples were then washed with PBS by centrifugation. The fixation buffer was added, and samples were incubated on ice for 30 min, followed by washing twice at 4°C with PBS by sedimentation and resuspension. Finally, cells were suspended in PBS and kept at 4°C until imaging. For dual-color co-localization analysis, two different proteins were labeled sequentially, using the same protocol described above. For imaging studies on cells transfected with PTPα-HA, 3HA-PM or GG-3HA, the same procedure was followed as described above, except that Alexa-647 tagged anti-HA antibody was used to label the expressed protein and probe.

For live cell imaging, IgE-sensitized cells were stained with FM143 dye (5 µg/mL) for 20 min at 4°C. Then cells were washed twice with PBS by centrifugation at 4°C, suspended in PBS at 4°C, and used within 15 min of completing the labeling process. For imaging, the cells were incubated at 37°C for 5 min just before dropping on antigen-activating or non-activating surfaces, prepared as described below.

For flow cytometry studies, suspended, non-sensitized or IgE-sensitized cells in resting or antigen-activated states were fixed as described above then incubated with selected antibody (specific for ERM, P-ERM, cofilin or P-cofilin) in the permeabilization buffer overnight at 4°C. Then the cells were washed twice with PBS and incubated with Alexa-568 labeled secondary antibody in the permeabilization buffer overnight at 4°C. The cells were then washed with PBS, and this buffer was exchanged with FACS buffer (PBS, 5mM EDTA, 1%BSA). These samples were immediately analyzed using a Thermo Fisher Attune NxT flow cytometer.

**1.6. Sample preparation for microscopy:** The cover-glass surfaces of MatTek dishes were cleaned by treating with 1 M NaOH for 40 min, then washed with distilled water and PBS several times. For fixed cell imaging, the cleaned MatTek dishes were coated with poly-D-lysine (PDL) for 40 min and washed with PBS. A small volume of the labeled cell suspension (~100 µl) in PBS was placed on the cover-glass region, and cells were allowed to settle on the PDL surface for 10 min. Then the PBS buffer was discarded and freshly prepared "blinking buffer" (50 mM cysteamine, 0.5 mg/mL glucose oxidase, 40 µg/mL catalase, 10% (wt/vol) glucose, 93 mM Tris·HCl, PBS buffer (pH 7.5–8.5)) was added. Imaging was conducted after 30 min.

For live cell imaging on activating or non-activating surfaces, the cleaned MatTek dishes were coated with either BSA solution (10 µg/ml, non-activating) or DNP-BSA (20 mol%)/BSA (total 10 µg/ml, activating) in Earles Balanced Salt Solution for 2 hr at room temperature. Then dishes were washed twice with PBS and immediately used for imaging with cells.

**1.7. TIRF setup:** Our microscope setup is described in detail in previous publications (3, 13). Briefly, the custom-built total internal reflection fluorescence microscopy setup is equipped with an inverted microscope (Leica DM-IRB, Wetzlar, Germany), with an oil immersion objective (PlanApo, 100´, NA 1.47; Leica Microsystems, Germany), a 488 nm excitation laser (Coherent, Santa Clara, CA), a 647 nm diode-pumped solid-state (DPSS) laser (Crystalaser, Reno, NV), an electron multiplying charge-coupled device (EMCCD) camera (back-illuminated Andor iXON Ultra 897, Andor Technology, Belfast, UK), OptoSplit II emission image splitter (Cairn Research, UK) and a piezo stage (PI nano Z-Piezo slide scanner; PI). The laser power is attenuated with neutral density filters as needed for the illumination requirements of the experiment. Free-space combination of the two laser lines was performed. The excitation laser beams were relayed through a pair of tilting mirrors and a dichroic mirror (ZT405/488/561/640rpc, Chroma Technology) and focused on the back focal plane of the objective. The EMCCD camera was employed to record the fluorescence signal after it passed through the same objective and the dichroic mirror and was spectrally separated by the OptoSplit II emission image splitter (with T560lpxrxt-UF2, ET575LP,

ET525/50M, or ZET 642nf filters, all from Chroma). The single EMCCD camera was utilized in a dual-view mode: Spectrally separated images were projected onto the two halves of the CCD chip.

**1.8. Microvillar Cartography (MVC) Analysis:**

**1.8.1. Reconstruction of 3D cell surfaces and detection of microvilli:** Detailed methodology for reconstruction of the 3D membrane surface and detection of microvilli from TIRFM images is described in previous publications (10, 12). In brief, a series of TIRF images were recorded of FM143- or FM143fx-labeled cells using different angles of incidence under weak illumination of a 488-nm laser. The closest point (pixel) to the glass surface should have maximum intensity for a particular angle of incidence ($I_{max}(\theta)$). We employed the following equation to compute the relative axial distance ( $\delta z$ ) of each point on the cell surface:

$$\delta z = \ln\left(I_{\max}(\theta)/I(\theta)\right)/d(\theta), \tag{1}$$

where $d(\theta)$ denotes penetration depth of the TIRF evanescent field with an angle of incidence θ, which is determined from the following equation:

$$d(\theta) = (\lambda/4\pi)\left(\eta_1^2 \sin^2\theta - \eta_2^2\right). \tag{2}$$

In this equation λ is the excitation wavelength (488 nm), and $\eta_1$ and $\eta_2$ are the refractive indices of the immersion oil (1.52) and blinking buffer (1.35), respectively.

A custom-built algorithm in MATLAB was developed to define the position of the tips of the microvilli. Initially, the TIRF images were processed with the “Laplacian of a Gaussian” (LoG) filter and binary images generated from the processed images by setting a threshold. Then MATLAB built-in function “bwlabel” was used to segment the binary images. The inclusion of the maximum number of microvilli was ensured by considering the binary image acquired from the data recorded at the smallest angle of incidence for segmentation first, and then data from the image recorded at the largest angle of incidence were combined to separate individual microvilli (Fig. S1D-H). Finally, segmented areas acquired from images taken at different areas were merged, and the mean of the δz of each pixel was determined and employed to generate the LocTips map. For a more detailed description, the reader is referred to our previous publication (See ref (10) - Section 4 – Notes - Point 23: ‘Analysis of VA-TIRFM images of cell membrane to reconstruct the 3-D topography of cell membrane’).

**1.8.2. Quantitative estimation of change of membrane topography:**

To quantify membrane topography we define the parameter of percentage microvillar (MV) area as compared to total plasma membrane area. For that purpose, the LocTips map (Fig. S1I-J) was used to define the projected microvillar area. The rest of the imaged area was divided into cell-body (CB; δz < 400 nm) and background (BG; δz > 400 nm) based on their distance from the glass surface. The total cell area is calculated by summing the MV and CB areas. Then the percentage of total MV area is calculated with respect to total cell area. These calculations are carried out for at least 10 cells for each condition of membrane topography evaluated, and the population means are compared with one way ANOVA test to determine whether they are significantly different at the P-value level of 0.05.

**1.8.3. Reconstruction of 3D membrane topography movie:** To map the membrane topography change of IgE-sensitized cells during interaction with activating or non-activating surfaces, we recorded the TIRF movie (frame time 100 ms) of live FM143-labeled cells during the interaction with those surfaces at a constant laser beam incident angle of 66.6°. Then the TIRF movie was divided into a series of 10 or 20 consecutive frames (i.e. 2 or 1 s, respectively) per segment. These 10 or 20 frames of each segment were merged, and the 3D cell surface reconstruction analysis described in the previous section was performed on each segment. From this procedure we obtained the membrane topography of the mast cell as a function of the interaction time with the surfaces. Then we clubbed the time-resolved membrane topography map of the cell to generate movies of 3D membrane topography changes of the cell during the interaction with activating and non-activating surfaces (SI Appendix Movies S1 and S2).

**1.8.4. Stochastic Localization Nanoscopy (SLN) and analysis:** We used Alexa-647 and Alexa-488 tags, which can be converted to a dark state upon excitation with a very high-intensity laser of the appropriate wavelength. The presence of the blinking buffer containing cysteamine together with glucose oxidase/catalase (an oxygen-scavenging system (14, 15)) ensures the switching back of a small fraction of the dye molecules at any moment of time. Thus, separate appearance of individual fluorophores is observed in consecutive recording frames, yielding precision of molecular localization well below the diffraction limit. For all SLN measurements, the laser beam incident angle was 66.6°. For each focal plane, we recorded 32,000 frames, in four series of movies (each containing 8000 frames). A piezo stage (PI nano Z-Piezo slide scanner) was employed to record SLN movies at the 0 nm and 400 nm focal planes. Our measurement at these two different slices guaranteed that proteins localized on membrane regions situated somewhat farther from the surface are not overlooked due to the length of the membrane projections. Moreover, considering the depth of focus and the high laser power used in our SLN measurements, focusing on the 400 nm slice enables visualization of fluorophores in cell body regions, as well as on microvilli surfaces. Detailed molecular localization analysis was published previously (ref (10) -

Section 4 – Notes - Point 22: 'Analysis of SLN movies to generate super-resolved map of membrane proteins'(10)).

**1.8.5. Drift correction:** TIRF reference images of the cell membrane were registered before and after each SLN movie under weak illumination of a 488-nm laser. A 2D cross-correlation of each reference TIRF image with the previous reference image was computed to identify and correct for any drift. We utilize the leak from 488 nm channel to the 647 nm channel to determine any drift between them. The leaked image in the 647 nm channel is cross-correlated with the original 488 nm image to identify and correct for any shift between the two channels' images. For dual-color SLN, we used fiducial markers (0.1 μm fluorescent microspheres, Tetraspeck) for drift correction.

**1.8.6. Quantitative analysis of membrane protein distributions:** We performed the segmentation of the membrane area of each cell into microvilli (MV) regions and non-microvilli or cell-body (CB) regions following the analysis described above (10, 12). The percentage of specifically labeled proteins in microvilli regions and on cell body regions were calculated for each cell (Fig. S1 I-J). Because we calculate the ratio for each protein individually in MV and CB regions, the fluorophore blinking rate does not affect our conclusion. We also calculated the density of specifically labeled proteins on each cell as a function of the distance from the central region of each microvillus. We defined the central microvillar region as that region within 20 nm from the microvillar tip (the pixel with the minimum δz value). A 'boundary' function in MATLAB was employed to encapsulate that central region, which may take different shapes depending on the specific microvilli and their alignment with respect to the membrane surface (Fig. S1L). Then, concentric closed curves of similar shapes in increasing increments of 20 nm were drawn. The number of fluorophores in each of these concentric regions was determined (Fig. S1M-N), and the density was calculated by dividing by the area of the respective regions. The resulting density values, δCount/δArea, *vs* distance (R) from the microvilli tip were plotted in units of 20nm. A steep slope of this δCount/δArea *vs* R plot indicates a rapid decrease in labeled proteins as the distance from the central microvillar region increases.

To determine the percentage of microvilli occupied by a specifically labeled protein we calculated the total number of microvilli from the segmentation map for each cell. Then we determined how many microvilli of that cell are occupied by at least one of the selected proteins and the corresponding percentage of the total.

**1.9. Quantitative estimation of co-localization level of membrane proteins:** We followed the co-localization probability (CP) analysis scheme developed previously (10, 12). This applies to cases where two labeled proteins are located on a single microvillus, and consequently in proximity, even if there is no interaction between them. This analysis depends on computing distances between localized points in super-resolved images. Briefly, the co-localization percentage of proteins $a$ and $b$ as $\frac{N_{ab}}{N_a} \times 100$, where $N_a$ is the total number of detected $a$ proteins, and $N_{ab}$ is the number of $a$ proteins that have at least one $b$ protein within the specified distance. The protein with the lower labeling density is always selected as the $b$ protein.

##### 1.10. TIRFM Imaging of Spatial Distribution of Signaling Proteins

The cells were labeled using the same procedure described above. TIRF movie (frame time 15 ms; Number of frames: 20 frames) of alexa-647 tagged antibody labeed fixed cells were recorded using a constant laser beam of 647 nm laser (incident angle of 66.6°). These 20 frames of each movie were merged to develop the TIRFM images.

##### 1.11. SEM Imaging:

All samples were fixed in 2% glutaraldehyde in 0.05M cacodylate buffer for 2 hr at 4°C. After the primary fix the cells were rinsed three times for 10 min at 4°C in 0.05M cacodylate buffer. The secondary fix was 1% osmium tetroxide in 0.05M cacodylate buffer, with incubation for 1 hr at 4°C under the hood. The osmium tetroxide was rinsed out with 0.05M cacodylate buffer three times for 10 min at 4°C. The cells were then distributed on Whatman #5 filter paper using a vacuum filter apparatus. The filter paper with the distributed cells were immediately transferred to a 9 cm plastic Petri dish containing 25% ethanol at 4°C for 10 min. The cells on filter paper were transferred to 50%-70%-95%-100%-100% ethanol at 4°C for 10 min each. We used a modified critical point drying process in which we let the samples soak for 24 hours in bone dry liquid carbon dioxide after the ethanol is rinsed out of the cells, ensuring that no ethanol or any other liquid remains in the cells, which can cause shrinkage in the cells as we heated the samples through the transition phase of the carbon dioxide. After the liquid carbon dioxide had gone through critical point, the cells were brought to atmospheric pressure over 30 min. The cells on filter paper were then mounted on an aluminum SEM mount using carbon conductive double stick tape and supplemented with a bead of silver paint from filter paper to mount to ensure conductivity. The mounted samples were then sputter coated with 15-20 nm of gold palladium and imaged at 1.75 kV with a KECK Leo 1550 SEM.

**2. Experiments to monitor changes in phosphorylation of cytoskeletal attachment proteins that accompany RBL mast cell activation.** We utilized flow cytometry and immunoblots to build on previous reports suggesting that the interplay between the ERM proteins and cofilin controls membrane topographical changes upon activation of T cells and B cells (12, 16, 17). Active forms of ERM proteins, i.e. phosphorylated ERM (P-ERM), are shown to stabilize the actin cytoskeleton and thereby support the formation of microvilli (12, 16, 17). The effect of ERM proteins is counterbalanced by cofilin proteins which when dephosphorylated increase actin severing, thereby alterring the formation of these membrane protrusions (17). We used antibodies specific for either phosphorylated forms or all forms of ERM and cofilin to label the endogenous ERM and cofilin proteins in RBL mast cells. After tagging with fluorescent secondary antibodies, we used flow cytometry to measure the amount of phosphorylated and total ERM and cofilin proteins in RBL cells in the presence and absence of C2-ceramide under four cell conditions: 1) resting cells not sensitized with IgE (NS), 2) IgE-sensitized cells with no (resting, S), or 3) 1 min, or 4) 15 min antigen activation (Fig. S5A-H). Using the NS cells as a normalizing baseline we calculated the percentage increase shift in the Mean Fluorescence Intensity (MFI) for ERM, P-ERM, cofilin and P-cofilin in the presence and absence of C2-ceramides after antigen activation (Fig. S5I-J).

MFI = % change of 'A' protein in 'X' condition, as compared to NS condition

$$= \frac{(MFI_X^A - MFI_{NS}^A)}{MFI_{NS}^A} \times 100$$

In the absence of C2-ceramide, P-ERM content of the cells appears to increase after antigen-activation for 1 min (57±16%) and 15 min (75±25%). A smaller increasing trend is also seen with total ERM content following antigen-activation for 1 min (32±11%), and 15 min (62±7%) (Fig. S5I). Correspondingly, the ratio of P-ERM (active) vs total ERM increases significantly after a minute of antigen activation (Fig. S5K), and this may be related to the initial merger of microvilli. The ratio of P-ERM to ERM decreases somewhat after 15 min (Fig. S5K) when most of the microvilli have merged into ruffles. Notably, in presence of C2-ceramide neither the P-ERM nor the ERM content increases significantly, after 1 min or 15 min of antigen addition (Fig. S5I). Considering that C2-ceramide causes a dramatic decrease in FcεRI/Lyn proximity (main text, Fig, 4) as well as disrupts signaling activity (18), these results are consistent with the view that effective antigen-stimulated coupling of FcεRI with Lyn increases ERM phosphorylation and merging of microvilli.

The flow cytometric results are supported qualitatively by visualization of immunoblots (Fig. S5M-P). The presence of two ERM bands in these blots is consistent with the expression of Moesin and Ezrin but not Radixin in mast cells (19). (We cannot rule out two bands for P-ERM.) The increase in total ERM protein expression we observe within minutes may be due to stimulated synthesis of this protein, known to be possible as early as 20 sec (20, 21). Our immunoblots also show an increase in the intensity of dual band for total ERM proteins within 1 minute of antigen activation compared to non-stimulated (NS) or antibody-sensitized (S) state of mast cells (Fig. S5M). Another possible contribution to the increase is a stimulated change in the accessibility of the labeling antibodies, which we expect to be modulated in the same way for anti-ERM and anti-P-ERM.

In contrast to the ERM case, phosphorylated cofilin and total cofilin do not appear to significantly increase after 1 min antigen activation (Fig. S5J). Both forms of the protein increase after 15 min antigen activation, with the increase of P-cofilin (26±8%) about half that of total cofilin (50±15%) (Fig. S5J). Overall, the changes of cofilin after antigen-activation are considerably less compared to ERM proteins as quantified by flow cytometry (Fig. S5I) and visualized in immunoblots (Fig. S5O and P). Because phosphorylation of cofilin inactivates this enzyme, the ratio of active cofilin *vs* total cofilin (Fig S5L) appears to increase after longer activation, corresponding to more severing of the actin cytoskeleton by cofilin. This may play a role in transforming microvilli by severing initial cytoskeletal attachments and merging these structures into ruffles. C2-ceramide treatment does not appear to have a significant effect on the expression of P-cofilin or total cofilin. Together our results indicate that FcεRI-mediated stimulation modulate the phosphorylation state of ERM proteins and less so for cofilin proteins.

**References:**


1. A. J. Torres, L. Vasudevan, D. Holowka, B. A. Baird, Focal adhesion proteins connect IgE receptors to the cytoskeleton as revealed by micropatterned ligand arrays. *Proc Natl Acad Sci U S A* **105**, 17238–44 (2008).
2. S. L. Veatch, *et al.*, Correlation Functions Quantify Super-Resolution Images and Estimate Apparent Clustering Due to Over-Counting. *PLoS One* **7**, e31457 (2012).
3. N. Bag, *et al.*, Lipid-based and protein-based interactions synergize transmembrane signaling stimulated by antigen clustering of IgE receptors. *Proceedings of the National Academy of Sciences* **118** (2021).
4. D. C. Seldin, *et al.*, Homology of the rat basophilic leukemia cell and the rat mucosal mast cell. *Proceedings of the National Academy of Sciences* **82**, 3871–3875 (1985).
5. E. L. Barsumian, C. Isersky, M. G. Petrino, R. P. Siraganian, IgE-induced histamine release from rat basophilic leukemia cell lines: isolation of releasing and nonreleasing clones. *Eur J Immunol* **11**, 317–323 (1981).
6. R. M. Z. G. Naal, J. Tabb, D. Holowka, B. Baird, In situ measurement of degranulation as a biosensor based on RBL-2H3 mast cells. *Biosens Bioelectron* **20**, 791–796 (2004).
7. M. D. Sharp, K. Pogliano, An *in vivo* membrane fusion assay implicates SpoIIIE in the final stages of engulfment during *Bacillus subtilis* sporulation. *Proceedings of the National Academy of Sciences* **96**, 14553–14558 (1999).

8. R. Rea, *et al.*, Streamlined Synaptic Vesicle Cycle in Cone Photoreceptor Terminals. *Neuron* **41**, 755–766 (2004).

9. K. H. R. Jensen, R. W. Berg, CLARITY-compatible lipophilic dyes for electrode marking and neuronal tracing. *Sci Rep* **6**, 32674 (2016).

10. S. Ghosh, A. Alcover, G. Haran, "Microvillar Cartography: A Super-Resolution Single-Molecule Imaging Method to Map the Positions of Membrane Proteins with Respect to Cellular Surface Topography" in *The Immune Synapse: Methods and Protocols*, C. T. Baldari, M. L. Dustin, Eds. (2023).

11. S. Ghosh, *et al.*, CCR7 signalosomes are preassembled on tips of lymphocyte microvilli in proximity to LFA-1. *Biophys J* **120**, 4002–4012 (2021).

12. S. Ghosh, *et al.*, ERM-Dependent Assembly of T Cell Receptor Signaling and Co-stimulatory Molecules on Microvilli prior to Activation. *Cell Rep* **30**, 3434-3447.e6 (2020).

13. S. A. Shelby, D. Holowka, B. Baird, S. L. Veatch, Distinct stages of stimulated FcεRI receptor clustering and immobilization are identified through superresolution imaging. *Biophys J* **105**, 2343–54 (2013).

14. S. van de Linde, *et al.*, Direct stochastic optical reconstruction microscopy with standard fluorescent probes. *Nat Protoc* **6**, 991–1009 (2011).

15. G. T. Dempsey, *et al.*, Photoswitching mechanism of cyanine dyes. *J Am Chem Soc* **131**, 18192–3 (2009).

16. S. Faure, *et al.*, ERM proteins regulate cytoskeleton relaxation promoting T cell–APC conjugation. *Nat Immunol* **5**, 272–279 (2004).

17. A. Droubi, *et al.*, The inositol 5-phosphatase INPP5B regulates B cell receptor clustering and signaling. *Journal of Cell Biology* **221** (2022).

18. D. Holowka, K. Thanapuasuwan, B. Baird, Short chain ceramides disrupt immunoreceptor signaling by inhibiting segregation of Lo from Ld Plasma membrane components. *Biol Open* **7** (2018).

19. T. C. Theoharides, D. Kempuraj, Potential Role of Moesin in Regulating Mast Cell Secretion. *Int J Mol Sci* **24**, 12081 (2023).

20. B. Alberts, *et al.*, *Molecular Biology of the Cell*, 4th edition (Garland Science; 2002) Chapter 6, Page 375.

21. M. Shamir, Y. Bar-On, R. Phillips, R. Milo, SnapShot: Timescales in Cell Biology. *Cell* **164**, 1302-1302.e1 (2016).

Table S1: P-Values from Fig. 2M, N and SI Fig. S2Q for distribution of signaling proteins with respect to membrane topography of RBL mast cells in resting condition and after 1 min and 15 min antigen-activation.

| **Data Sets tested with one-way ANOVA** | **P Value** | | |
|---|---|---|---|
| | **% of Protein on MV** | | **% of MV with Protein** |
| | 0 nm | 400 nm | |
| FcεRI vs FcεRI+1min Ag | 0.31 | 0.38 | 0.09 |
| FcεRI vs FcεRI+15 min Ag | 0.84 | 0.88 | 0.07 |
| FcεRI+1min Ag vs FcεRI+15 min Ag | 0.38 | 0.10 | 0.67 |
| Lyn vs Lyn+1min Ag | 0.48 | 0.46 | 0.28 |
| Lyn vs Lyn+15 min Ag | 0.62 | 0.23 | 0.58 |
| Lyn+1min Ag vs Lyn+15 min Ag | 0.73 | 0.65 | 0.80 |
| LAT vs LAT+1min Ag | **$0.04 \times 10^{-2}$** | **0.02** | 0.82 |
| LAT vs LAT+15 min Ag | **$0.07 \times 10^{-5}$** | **$0.01 \times 10^{-5}$** | 0.85 |
| LAT+1min Ag vs LAT+15 min Ag | **$0.03 \times 10^{-5}$** | **$0.01 \times 10^{-5}$** | 0.97 |
| PTPα vs PTPα+1min Ag | 0.11 | **0.02** | **0.01** |
| PTPα vs PTPα+15 min Ag | 0.77 | 0.86 | 0.19 |
| PTPα+1min Ag vs PTPα+15 min Ag | 0.24 | 0.09 | 0.13 |

Table S2: P-Values for distribution of probes preferring either ordered lipid (PM) or disordered lipid (GG) regions (Fig. 3E,F and SI Fig. S3E) and signaling proteins (Fig. 3Q,R and SI Fig, S3M) with respect to membrane topography of RBL mast cells in resting condition, with or without C2-ceramide treatment and continuously C2-Ceramide treated cells after 15 min antigen-activation.

| Data Sets tested with one-way ANOVA | P Value | | |
|---|---|---|---|
| | % of Protein on MV | | % of MV with Protein |
| | 0 nm | 400 nm | |
| PM vs PM+C2 | 0.32 | 0.36 | **0.07 x $10^{-7}$** |
| GG vs GG+C2 | 0.54 | 0.75 | 0.82 |
| FcεRI vs FcεRI+C2 | 0.94 | 0.30 | **0.03 x $10^{-2}$** |
| FcεRI vs FcεRI+C2+15 min Ag | 0.89 | 0.56 | **0.03 x $10^{-3}$** |
| FcεRI+C2 vs FcεRI+C2+15 min Ag | 0.85 | 0.14 | 0.97 |
| Lyn vs Lyn+C2 | 0.06 | 0.23 | **0.05 x $10^{-3}$** |
| Lyn vs Lyn+C2+15 min Ag | 0.30 | 0.37 | **0.03 x $10^{-3}$** |
| Lyn+C2 vs Lyn+C2+15 min Ag | **0.02 x $10^{-1}$** | 0.08 | 0.70 |
| LAT vs LAT+C2 | **0.02 x $10^{-4}$** | **0.08 x $10^{-3}$** | **0.01** |
| LAT vs LAT+C2+15 min Ag | **0.07 x $10^{-4}$** | **0.06 x $10^{-4}$** | 0.93 |
| LAT+C2 vs LAT+C2+15 min Ag | 0.45 | 0.76 | **0.02** |

Table S3: P-Values for relative degranulation (SI Figs. S2S and S3O)

| **Data Sets tested with one-way ANOVA** | **P Value** |
|---|---|
| Untransfected spontaneous vs Untransfected antigen-activated | **$0.02 \times 10^{-1}$** |
| PTPα transfected spontaneous vs PTPα antigen-activated | **$0.03 \times 10^{-1}$** |
| Untransfected spontaneous vs PTPα transfected spontaneous | 0.86 |
| Untransfected antigen-activated vs PTPα antigen-activated | **$0.03 \times 10^{-1}$** |
| Untreated spontaneous vs Untreated antigen-activated | **$0.06 \times 10^{-4}$** |
| C2-ceramide treated spontaneous vs C2-ceramide treated antigen-activated | 0.22 |
| Untreated spontaneous vs C2-ceramide treated spontaneous | 0.64 |
| Untreated antigen-activated vs C2-ceramide treated antigen-activated | **$0.03 \times 10^{-3}$** |

Table S4: P-Values for Percentage of Mean Fluorescent Intensity (MFI) difference of ERM / PERM, and Cofilin / P-Cofilin with respect to resting state in the presence or absence of C2-ceramide in RBL mast cells calculated from flow cytometric histogram (SI Fig. S5I, J).

| **Data Sets tested with one-way ANOVA** | **P Value** |
|---|---|
| PERM+1min Ag vs PERM+15 min Ag | 0.57 |
| ERM+1min Ag vs ERM+15 min Ag | **0.03** |
| PERM+C2+1min Ag vs PERM+C2+15 min Ag | 0.88 |
| ERM+C2+1min Ag vs ERM+C2+15 min Ag | 0.79 |
| PCOF+1min Ag vs PCOF+15 min Ag | **0.01** |
| COF+1min Ag vs COF+15 min Ag | **0.04** |
| PCOF+C2+1min Ag vs PCOF+C2+15 min Ag | 0.31 |
| COF+C2+1min Ag vs COF+C2+15 min Ag | **$0.06 \times 10^{-1}$** |

### Figure S1

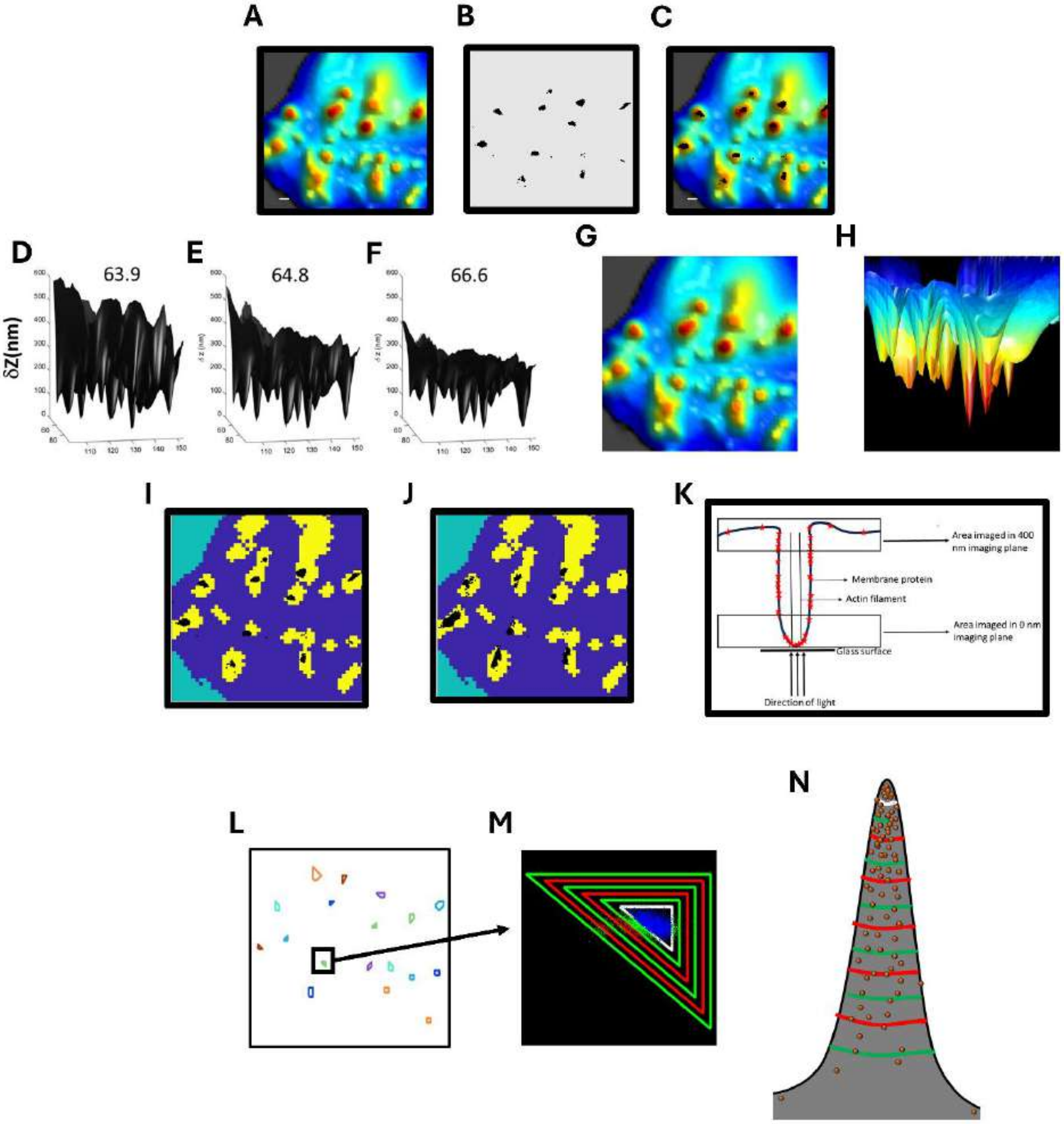
A
B
C
D
63.9
δZ(nm)
E
64.8
F
66.6
G
H
I
J
K
Area imaged in 400 nm imaging plane
Membrane protein
Actin filament
Area imaged in 0 nm imaging plane
Glass surface
Direction of light
L
M
N

**Figure S1. Schematic for Analysis of RBL Mast Cells with Microvillar Cartography**

A-C. Schematics of microvillar cartography (MVC) imaging. A. Representative membrane topography map obtained from variable angle total internal reflection fluorescence microscopic (VATIRFM) imaging. B. Representative localization map of membrane proteins (Lyn) obtained from stochastic localization nanoscopy (SLN). C. Merging of these two images enables localization of membrane proteins with respect to three-dimensional membrane topography.

D-F. Reconstruction of 3D surface of an RBL cell from VA-TIRFM and determination of microvilli regions. A series of 3D surface reconstruction maps of cell from TIRFM images, based on calculated δz values from TIRFM measurements at angles of incidence 63.9° (D), 64.8° (E), and 66.6° (F) under weak illumination of a 488-nm laser. The δz map is viewed perpendicular to the y–z plane.

G. Representative 3D topographical map of an RBL cell, reconstructed from the mean of δz values from D–F. The δz values are represented by different hues with a step size of 6.25 nm.

H. Bottom-to-top 3D view of G.

I.-J. Schematic of method for sorting membrane proteins into microvilli (MV, yellow) and cell-body (CB, blue) regions. Localizations of fluorescently labeled Lyn proteins (black dots) obtained with SLN from the 0 nm slice (I) and the 400 nm slice (J) are overlaid with the map that segments the two regions.

K. Schematic of observation regions in 0 nm and 400 nm imaging planes.

L.-N. Schematic for determining the fractional change in the number of blinking fluorophores labeling a specific protein on each RBL cell as a function of the distance (R) from the central region of each microvillus.

L. Descending microvillar regions to count the number of molecules as a function of the distance from the microvillar tip are marked with different colors.

M. A representative plot of the concentric shapes drawn around the central microvilli region (marked with the black box in L). The white closed curve in M is the central microvillar region.

N. Schematic of 3D representation of M (‘δCount/δArea’ analysis) with respect to a microvillar structure. White line represents the boundary of central microvillar region. Green and red lines are concentric shapes drawn around central microvillar region. Brown spheres represent the membrane proteins.

### Figure S2

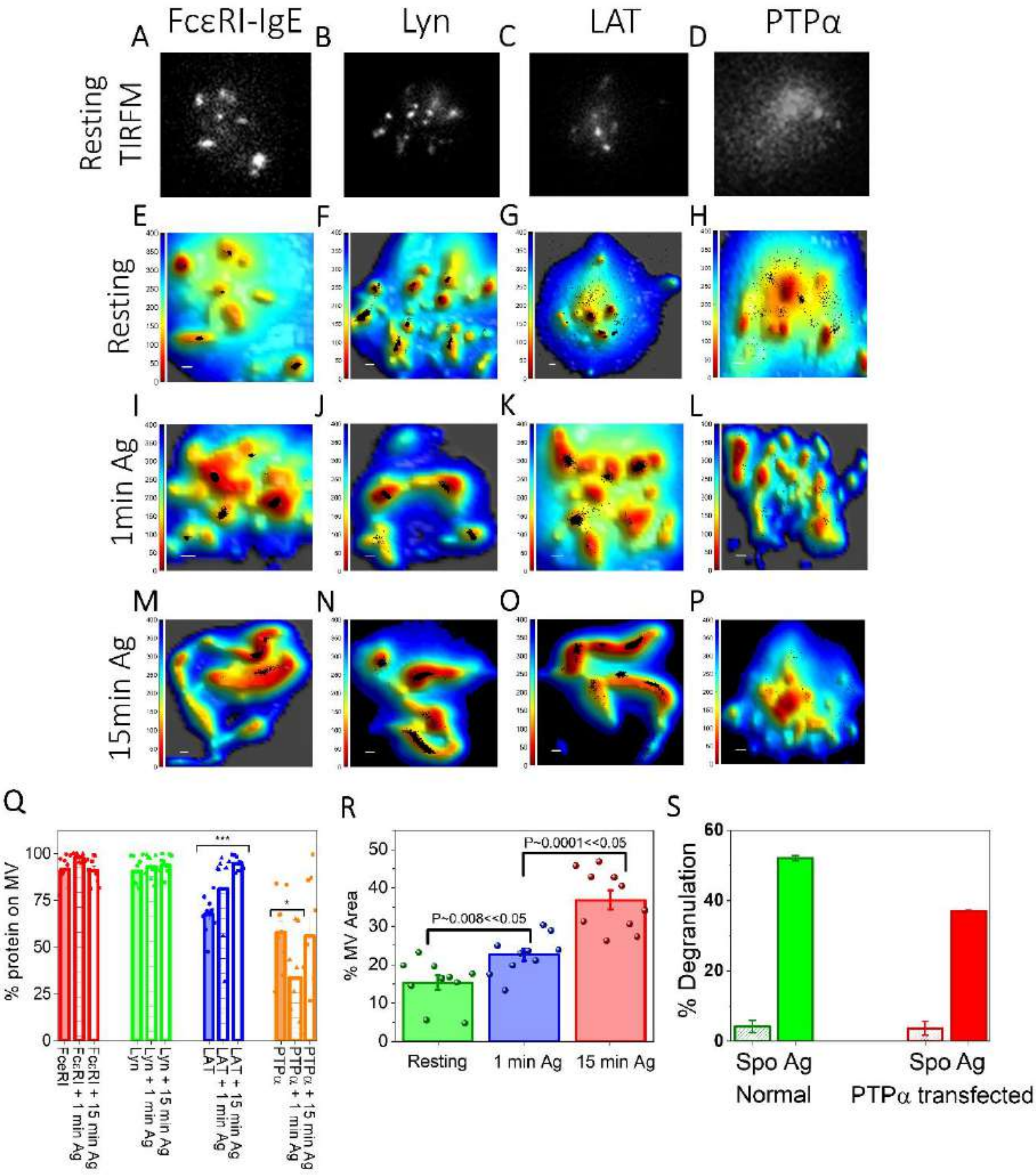

A FcεRI-IgE
B Lyn
C LAT
D PTPα
Resting TIRFM
E
F
G
H
Resting
I
J
K
L
1min Ag
M
N
O
P
15min Ag
Q
% protein on MV
FcεRI
FcεRI + 1 min Ag
FcεRI + 15 min Ag
Lyn
Lyn + 1 min Ag
Lyn + 15 min Ag
LAT
LAT + 1 min Ag
LAT + 15 min Ag
PTPα
PTPα + 1 min Ag
PTPα + 15 min Ag
R
% MV Area
P~0.008<<0.05
P~0.0001<<0.05
Resting
1 min Ag
15 min Ag
S
% Degranulation
Spo Ag
Normal
Spo Ag
PTPα transfected

**Figure S2. Membrane Topography and Localization of Signaling Proteins are Correlated in Resting and Antigen-Activated States of RBL Mast Cells:**

A-D. Representative TIRF images of RBL mast cells labeled with alexa-647 tagged antibody specific for FcεRI, Lyn, LAT, and PTPα, respectively in solution phase when the microvillar-dominated topography of mast cells are unaltered.

E-P. Localization maps of RBL cell signaling proteins with respect to the 3D membrane topography in suspended resting cells (E-H), and after 1 min (I-L) or 15 min (M-P) activation with soluble antigen (DNP-BSA). Positions of proteins obtained from SLN (black dots) in the 400 nm imaging slice are superimposed on membrane topography maps obtained from VA-TIRFM. The color bars represent the distance from the glass in nanometers (nm). Scale bars, 0.5 μm.

Q. Percentage of proteins on microvillar (MV) regions of the membrane in resting, 1 min, and 15 min antigen-activated states in the 400 nm imaging slice. The values for individual cells are shown as dots in the plot. Error bars represent standard error of mean.

R. Comparison of the percentage of microvillar area in suspended resting RBL mast cells, and after 1 min or 15 min activation with soluble antigen (DNP-BSA).

S. Degranulation of untransfected and PTPα transfected RBL cells as measured in standard plate assay. 'Spo' and 'Ag' correspond to spontaneous and antigen-activated cells. Error bar represents standard error of mean from a single experiment representing three repeated experiments.

P-values for S are given in SI Table S3. P-values for Q are given in SI Table S2

### Figure S3

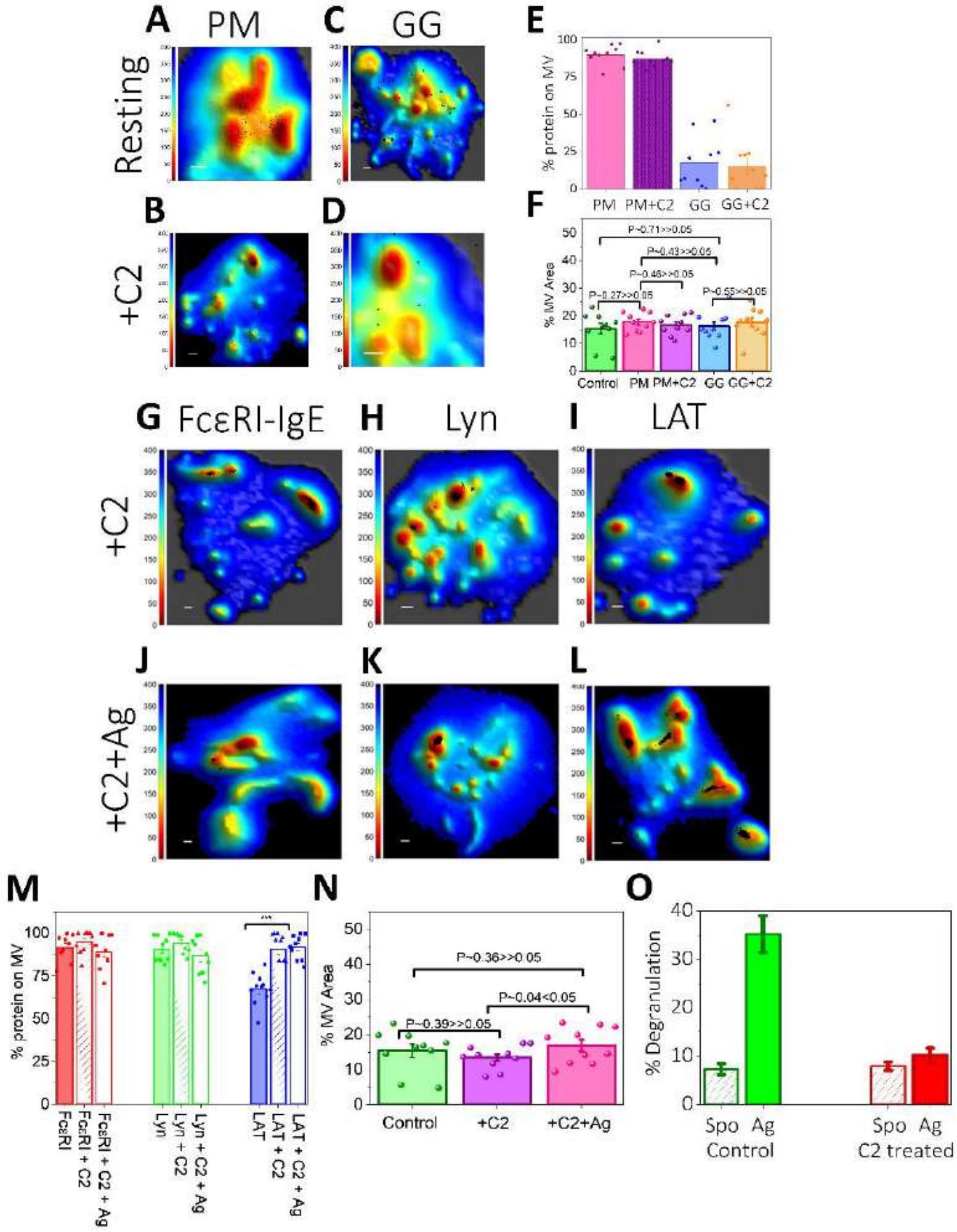

A PM
C GG
E
Resting
B
D
F
+C2
% protein on MV
PM PM+C2 GG GG+C2
% MV Area
P~0.71>>0.05
P~0.43>>0.05
P~0.46>>0.05
P~0.27>>0.05
P~0.55>>0.05
Control PM PM+C2 GG GG+C2
G FcεRI-IgE
H Lyn
I LAT
+C2
J
K
L
+C2+Ag
M
N
O
% protein on MV
FcεRI
FcεRI + C2
FcεRI + C2 + Ag
Lyn
Lyn + C2
Lyn + C2 + Ag
LAT
LAT + C2
LAT + C2 + Ag
% MV Area
P~0.36>>0.05
P~0.04<0.05
P~0.39>>0.05
Control
+C2
+C2+Ag
% Degranulation
Spo Ag
Control
Spo Ag
C2 treated

**Figure S3. Synergy between Membrane Topography and Membrane Domains Controls the Localization of Signaling Proteins in Resting and Antigen-Activated States of RBL Mast Cells:**

A-D. Localization maps of probes preferring either ordered lipid (PM-3HA; A-B) or disordered lipid (3HA-GG; C-D) regions with respect to 3D membrane topography in untreated (A, C), and C2-ceramide treated (B, D) resting cells. Positions of probes obtained from SLN (black dots) in the 400 nm imaging slice are superimposed on membrane topography maps obtained from VA-TIRFM. The color bars represent the distance from the glass in nanometers (nm). Scale bars, 0.5 μm.

E. Percentage of molecules on microvillar (MV) regions of the membrane in resting untreated and C2-ceramide treated suspended cells in the 400 nm imaging slice. The values for individual cells are shown as dots in the plot. The error bar represents the standard error of the mean.

F. Comparison of percentage of microvillar area in resting, PM-3HA transfected, C2-ceramide treated PM-3HA transfected, 3HA-GG transfected, and C2-ceramide treated 3HA-GG transfected RBL mast cells.

G-L. Localization maps of signaling proteins with respect to three-dimensional membrane topography in C2-ceramide treated (G-I) cells and after 15 min antigen-activation in continued presence of C2-ceramide (J-L). Positions of protein molecules obtained from SLN (black dots) in the 400 nm imaging slice are superimposed on membrane topography maps obtained from VA-TIRFM. The color bars represent the distance from the glass in nanometers (nm). Scale bars, 0.5 μm.

M. Percentage of proteins on microvillar (MV) regions of the membrane in the 400 nm imaging slice for untreated and C2-ceramide treated resting cells, and after 15 min antigen-activation in continued presence of C2 ceramide states. The values for individual cells are shown as dots in the plot. The error bar represents the standard error of the mean.

N. Comparison of percentage of microvillar area in resting, C2-ceramide treated resting, and after 15 min antigen-activation in the continued presence of C2 ceramide states of RBL mast cells.

O. Degranulation of untreated and C2-ceramide treated RBL cells as measured in the standard suspended cell assay. ‘Spo’ and ‘Ag’ correspond to spontaneous and antigen-activated cells. Error bar represents standard error of mean from a single experiment representing three repeated experiments.

P-values for E and M are given in SI Table S2. P-values for O are given in SI Table S3.

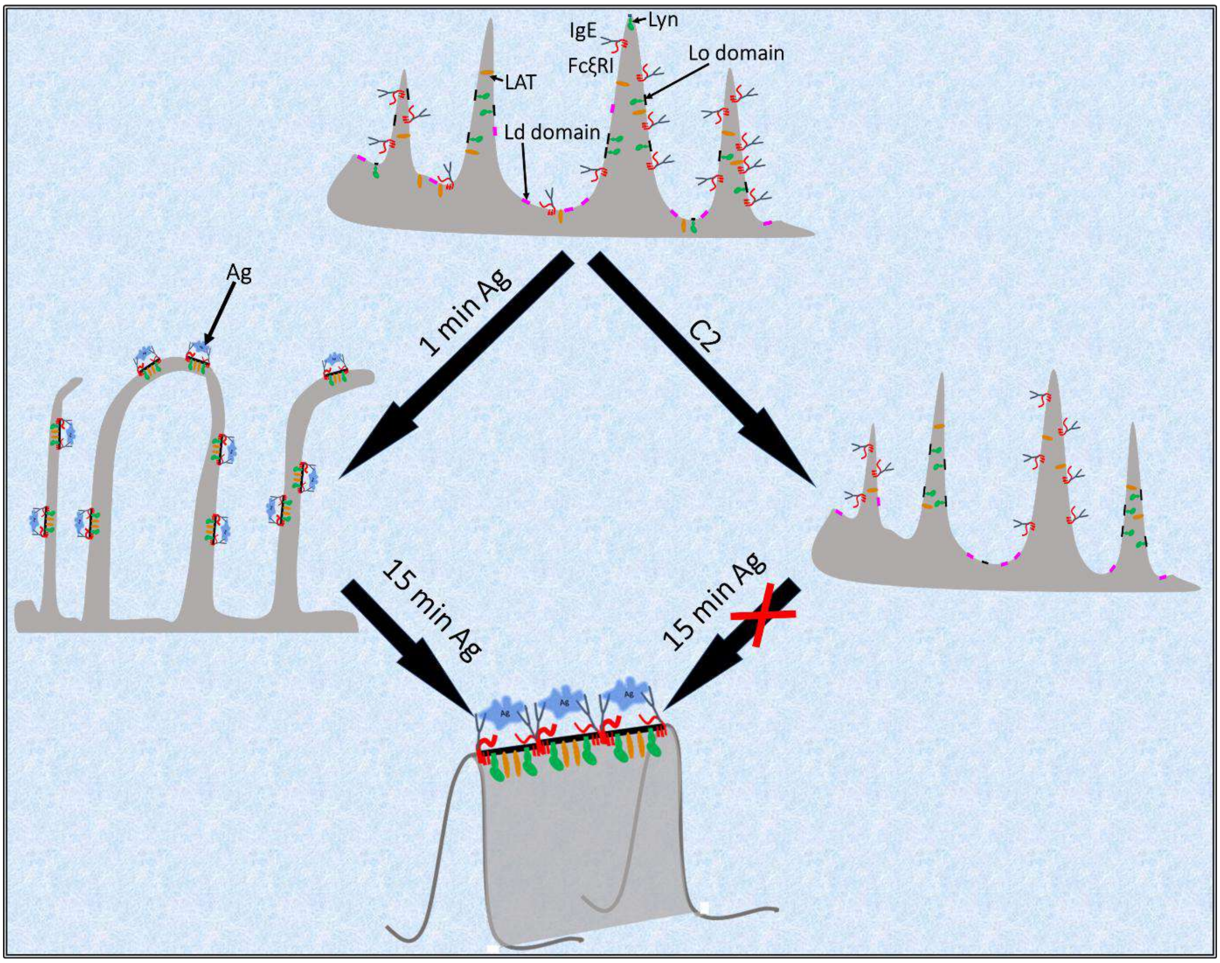


**Figure S4: Schematic of the Spatial Relationship between Topography and Lipid Domains in Plasma Membrane of RBL Mast Cells to Control the Localization of Signaling Proteins in Microvillar → Ruffle Projections and Facilitate Coordinated Activation after Antigen Engagement**

### Figure S5

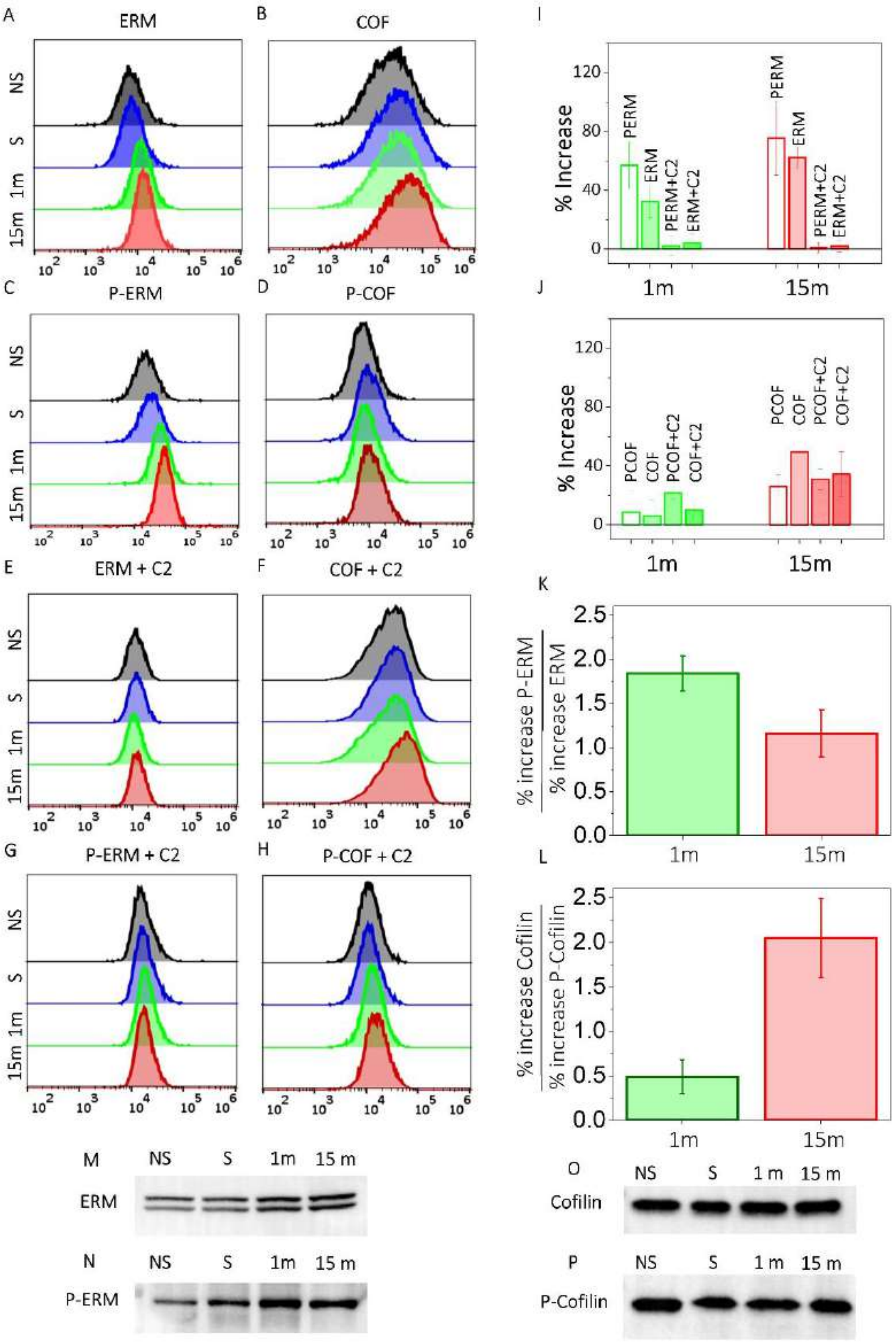
A
ERM
B
COF
C
P-ERM
D
P-COF
E
ERM + C2
F
COF + C2
G
P-ERM + C2
H
P-COF + C2
NS
S
1m
15m
I
% Increase
PERM
ERM
PERM+C2
ERM+C2
J
PCOF
COF
PCOF+C2
COF+C2
K
% increase P-ERM
% increase ERM
L
% increase Cofilin
% increase P-Cofilin
M
NS
S
1m
15 m
ERM
N
P-ERM
O
Cofilin
P
P-Cofilin

**Figure S5: Cytoskeleton Attachment Proteins Participate in RBL Mast Cell Activation:**

A-H. Comparison of flow cytometric analysis histograms of ERM (A), Cofilin (B), P-ERM (C), P-Cofilin (D), ERM in presence of C2-ceramide (E), Cofilin in presence of C2-ceramide (F), P-ERM in presence of C2-ceramide (G) and P-Cofilin in presence of C2-ceramide (H) in resting RBL cells not sensitized with IgE (NS), IgE-sensitized resting cells (S), 1 minute after antigen-activation (1m) and 15 minutes after activation (15m).

I-J. Percentage of Mean Fluorescent Intensity (MFI) difference of ERM-PERM (I), and Cofilin-P-Cofilin (J) with respect to non-sensitized resting state in the presence or absence of C2-ceramide in RBL cells, as calculated from flow cytometric histogram. The error bar represents standard error of the mean from three independent experiments.

MFI = % change of ‘A’ protein in ‘X’ condition, as compared to NS condition, as given by

$$\frac{(MFI_X^A - MFI_{NS}^A)}{MFI_{NS}^A} \times 100$$

K. Ratio of percentage of increase of P-ERM (active) vs total ERM after 1 minute (1m) and 15 minutes (15m) of antigen-activation. The error bar represents standard error of the mean from three independent experiments.

L. Ratio of percentage of increase of Cofilin vs P-Cofilin (inactive) after 1 minute (1m) and 15 minutes (15m) of antigen-activation. The error bar represents standard error of the mean from three independent experiments.

M-P. Immunoblot with lysates from resting RBL cells not sensitized with IgE (NS), IgE-sensitized resting cells (S), 1 minute after antigen-activation (1m) and 15 minutes after activation (15m). SDS-PAGE gels of cell lysates blotted with anti-ERM (M), anti-P-ERM (N), anti-Cofilin (O), or anti-P-cofilin (P) antibody. The complete western blots are shown in SI Fig. S6.

P-values for I and J are given in SI Table S4.

### Figure S6

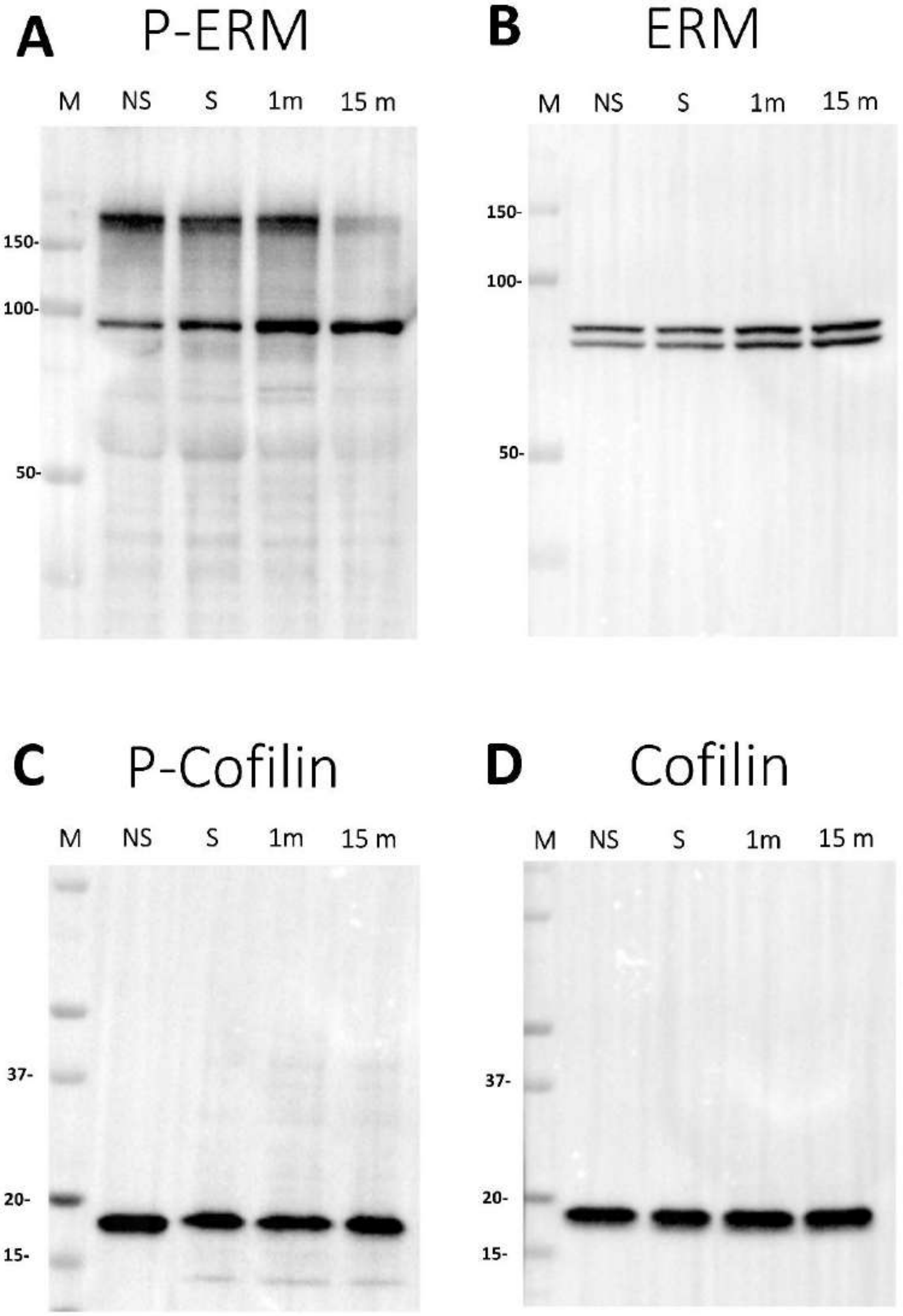

A
P-ERM
M NS S 1m 15 m
150-
100-
50-
B
ERM
M NS S 1m 15 m
150-
100-
50-
C
P-Cofilin
M NS S 1m 15 m
37-
20-
15-
D
Cofilin
M NS S 1m 15 m
37-
20-
15-

**Figure S6: Cytoskeleton Attachment Proteins Participate in RBL Mast Cell Activation:**
A-D. Complete Immunoblots corresponding to those shown in SI Fig. S5, with lysates from resting cells not sensitized with IgE (NS), IgE-sensitized resting cells with (S), 1 minute after antigen-activation (1m) and 15 minutes after activation (15m). SDS-PAGE gel of cell lysates blotted with anti-P-ERM (A), anti- ERM (B), anti-P-Cofilin (C), or anti-cofilin (D) antibody. M indicates the lane for molecular weight markers in kDa. The band above 150 kD in A is due to nonspecific binding of anti-P-ERM.

**Movie S1: Interaction of IgE-sensitized membrane labeled (FM143) RBL mast cell with only BSA coated non activating surface:**

A. 3D surface topographical map of an RBL cell, reconstructed from the mean of δz values as represented by different hues with a step size of 6.25 nm.
B. Bottom-to-top 3D view of A.

**Movie S2: Interaction of IgE-sensitized membrane labeled (FM143) RBL mast cell with 20 mol% DNP-BSA/BSA coated activating surface:**

A. 3D surface topographical map of an RBL cell, reconstructed from the mean of δz values as represented by different hues with a step size of 6.25 nm.
B. Bottom-to-top 3D view of A.
